# Control of NK cell tolerance in MHC class I-deficiency by regulated SHP-1 localization to the activating immune synapse

**DOI:** 10.1101/2022.03.08.483415

**Authors:** Laurent Schmied, Thuy T. Luu, Jonas N. Søndergaard, Stephan Meinke, Dara K. Mohammad, Sunitha B. Singh, Corinna Mayer, Giovanna P. Casoni, Michael Chrobok, Heinrich Schlums, Giorgia Rota, Hieu M. Truong, Lisa S. Westerberg, Greta Guarda, Evren Alici, Arnika K. Wagner, Nadir Kadri, Yenan T. Bryceson, Mezida B. Saeed, Petter Höglund

## Abstract

Signaling via inhibitory KIR/Ly49 receptors preserves natural killer (NK) cell self-tolerance but also conveys NK cell reactivity towards MHC class-I low target cells in an education process. Here, we demonstrate that mouse NK cell education by H-2D^d^ regulates transcription of several genes in Ly49A+ NK cells including *Ptpn6*, encoding the phosphatase SHP-1. SHP-1 was highly expressed in uneducated NK cells, in which knock-out of *Ptpn6* increased responsiveness. Following NKp46 triggering of uneducated NK cells, a higher synaptic abundance of phosphorylated SHP-1 was found relative to educated NK cells, concomitant with reduced phosphorylation of several signaling molecules, including PLC-g2, SLP-76, ZAP70/Syk and ERK1/2. SHP-1 overlapped extensively with F-actin and SLP-76 in the uneducated activating synapse of Ly49A+ NK cells, whereas a greater association between Ly49A and SHP-1 was observed in educated NK cells. Thus, our results indicate that in addition to transcriptional regulation, a distinct SHP-1 patterning in NK cell activating synapses can determine their tolerance.

## INTRODUCTION

Natural killer (NK) cells are innate lymphoid cells considered especially important in antiviral immune responses and tumor surveillance. Apart from direct cytotoxicity, NK cells produce proinflammatory cytokines such as IFN-γ, to modulate the activity of other immune cells, including those of the adaptive immune system ^1, 2^. NK cell activation is tightly controlled by germline-encoded activating and inhibitory receptors ^1, 3, 4^. Activating receptors (AR) recognize microbial or stress-associated self-molecules, expressed at low abundance in normal physiological condition but increased upon microbial infections or cellular stress ^5^. AR either have cytoplasmic immune-receptor tyrosine-based activation motifs (ITAM) themselves or associate with adaptor proteins that contain such motifs. These ITAMs are phosphorylated upon activation, recruiting SYK family kinases that transmit signals leading to an increase in intracellular Ca^2+^ and actin-dependent cytoskeletal rearrangement, which facilitates cell adhesion, synapse formation, degranulation and cytokine production ^1^.

Endogenous healthy cells resist NK cell killing due to their constitutive expression of self-MHC class I molecules, which bind inhibitory receptors (IR) on NK cells and thereby ensure self-tolerance. IR for MHC class I (MHC-I) are polymorphic and include C-type lectin homodimer receptors in mice (Ly49 family), killer immunoglobulin-like receptors (KIRs) in humans and C-type lectin heterodimer receptors (CD94-NKG2A) in both species ^6^. In addition to maintaining self-tolerance, IR promote the acquisition of NK cell function in a process termed “education” ^2, 7, 8, 9, 10^. The importance of this process was first demonstrated in mice expressing transgenic MHC-I molecules ^7, 11^ or lacking genes encoding β2-microglobulin, MHC-I heavy chain or transporter associated with antigen processing (TAP) ^9, 12, 13, 14^. Later, NK cells in TAP-deficient humans were also shown to be hyporesponsive ^15^.

Different NK education models have been proposed to account for the paradoxical “instructive” role of IR in NK cell responsiveness. The “arming” or “licensing” model proposes an education process in which inhibitory signals actively render NK cells functional ^16^. The “disarming” model, on the other hand, suggests that IR prevent overactivation and anergy which would otherwise result from unopposed activating signals triggered by the constitutive expression of activating ligands on surrounding cells, including ligands for NKG2D, NKp46, NK1.1 or SLAM family receptors ^17, 18, 19^. The “rheostat” model, which is compatible with both “arming” and “disarming”, is not an education model itself, but a way to explain the dynamic properties of NK cell education, i.e. how NK cells are fine-tuned by the level of IR and MHC-I molecules that they encounter in interactions with surrounding cells ^20, 21, 22, 23^. Despite the profound impact of NK cell education by MHC-I molecules, the molecular mechanisms underlying the process remain poorly understood.

IR signal immune-receptor tyrosine-based inhibitory motifs (ITIM) in their cytoplasmic tails. Binding of IR to MHC-I or MHC-I-like molecules induces ITIM phosphorylation, with subsequent recruitment and activation of tyrosine phosphatases, including Src homology region 2 domain (SH2)-containing phosphatase 1 (SHP-1), SHP-2 or SH2-containing inositol 5’ polyphosphatase 1 (SHIP1) ^6^. A recent study from Zhu and colleagues indicates SHP-1 expression levels and NK cell functional responsiveness are tightly linked ^24^, suggesting involvement in the molecular control of the rheostat that determines NK cell responsiveness ^21, 22, 23, 25^. The link between SHP-1 and NK cell maturation remains to be investigated, and it remains unclear how SHP-1 exert its role as a rheostat.

Here, we compared an MHC-I knock-out (KO) mouse strain (denoted as Dd−/− mice) and variants of this strain expressing either one or two alleles of H-2D^d^ transgene (denoted as Dd+/− and Dd+/+ mice) in which Ly49A+ NK cells are the most responsive subset ^7, 20^. Using this reductionistic model to study NK cell education, we uncover both transcriptional and post-transcriptional control of NK cell tolerance by SHP-1, the latter involving differential recruitment of SHP-1 to the immune synapse in uneducated *versus* educated NK cells following activation. Based on our observations, we postulate a model in which NK cell tolerance is controlled by SHP-1-mediated dephosphorylation of activating signaling molecules in the activating immune synapse. Following education by IR engagement, SHP-1 is sequestered from the immune synapse, releasing inhibition and potentiating NK cell responsiveness.

## RESULTS

### Dd expression in MHC-I KO mice affects NK cell single cell transcriptome

We hypothesized that the interaction between inhibitory Ly49 receptors and MHC class I induces key transcriptional changes for NK cell education. To study this, we set up a reductionistic NK cell education system in the C57Bl/6 (B6) mouse based on the comparison between MHC-I-deficient B6 mice and the same mice expressing either one or two copies of a single transgenic MHC-I gene, H2D^d 7, 20, 26^. Mice are denoted as Dd-/-, Dd+/− and Dd+/+ mice throughout the paper. We used flow cytometry to stratify cells into subsets expressing only one of the five main IR in B6 mice, including Ly49A, Ly49C, Ly49G2, NKG2A and Ly49I, and developed a boolean gating strategy to analyze and isolate them as single positive (sp) subsets **(Fig. S1a)**. First, we confirmed our previous data ^20, 21^ that all NK cells in Dd−/− mice are hypo-functional and that the introduction of either one or two copies of a single MHC-I transgene conveys quantitative education of sp-Ly49A NK cell **(Fig. 1a, b).** A difference in responsiveness was also seen for sp-Ly49C, sp-Ly49G2 and sp-NKG2A NK cells, although the difference for Ly49G2 did not reach statistical significance **(Fig. 1b)**. Among tested subsets, sp-Ly49I NK cells were the only subset not educated by H2D^d^ **(Fig. 1b)**. Thus, education by H2D^d^ according to the rheostat model is a general phenomenon for Ly49A, Ly49G2, Ly49C and NKG2A in H2D^d^ transgenic mice.

**Fig. 1.**
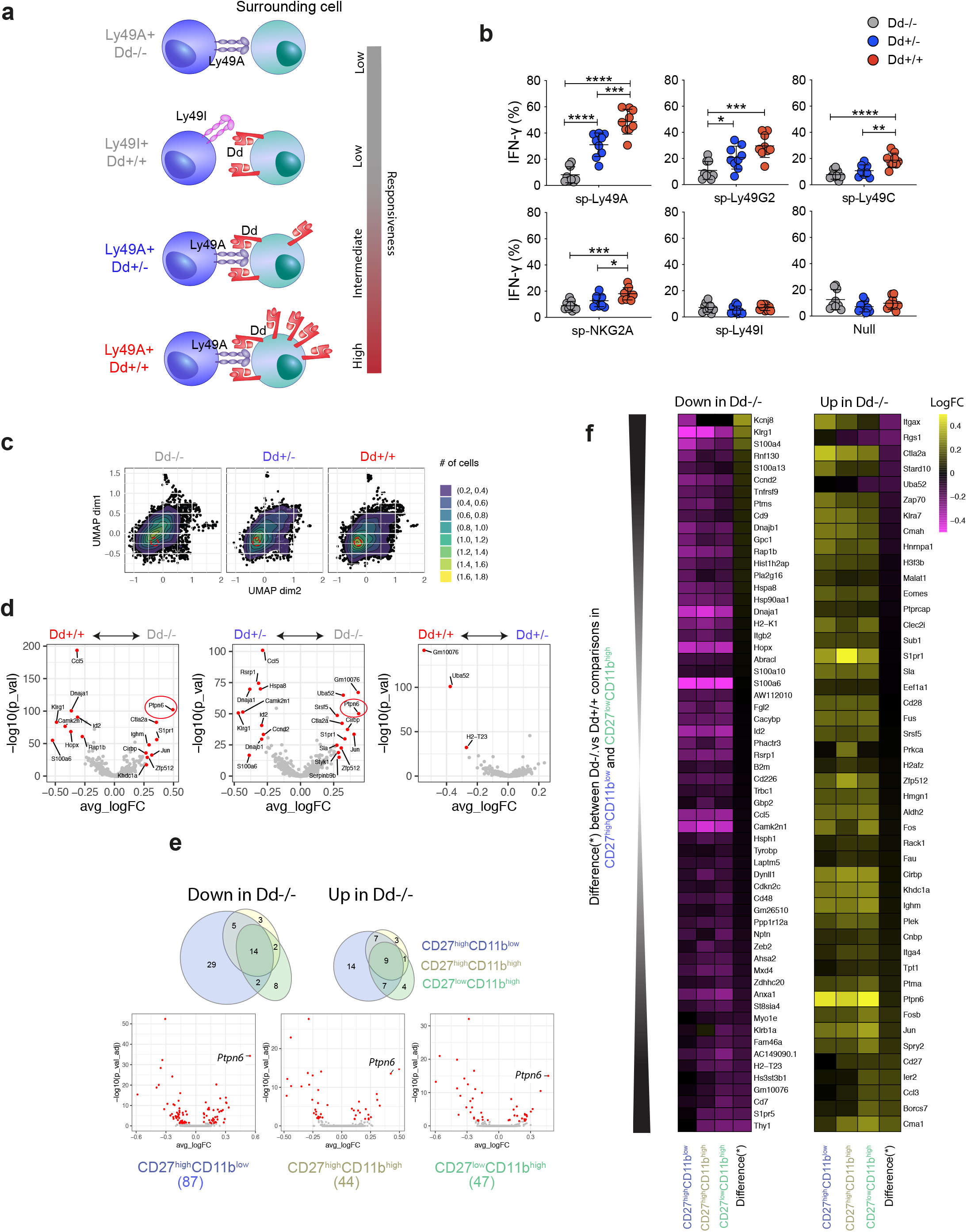
Single cell RNA-sequencing reveals transcriptional differences between educated and uneducated NK cells. **a,** Schematic model illustrating our mouse models. **b,** IFN-γ production in response to plate-bound NK1.1 crosslinking of NK cell subsets from Dd+/+, Dd+/− and Dd−/− mice. **c,** Uniform Manifold Approximation and Projection (UMAP) plot of single cell RNA sequencing (scRNAseq) data from sp-Ly49A subset (n=2 mice from each strain). Red dotted circles showed the overlay of the region with the highest density for Dd+/+ cells. **d,** Volcano plot showing DEGs. Red dots (FDR<0.05, FC>1.2) and grey dots (FDR>0.05). *Ptpn6*, is highlighted with a red circle. e, Venn diagram (top panel) and Volcano plots (bottom panel) showing the overlap of DEGs from three different maturation stages. Numbers below the Volcano plots indicated the numbers of DEGs in each comparison. **f,** Heatmap of the DEG genes from the comparisons in **e**.

Single cell RNA sequencing (scRNA-seq) on sorted sp-Ly49A NK cells from the three strains was performed. To minimize batch-effects, sorted NK cells from each strain were hash-tagged with different oligo-labelled antibodies recognizing all hematopoietic cells, pooled and sequenced together **(Fig. S1b-c)**. The initial quality control revealed an unspecific positive H2D^d^ signal in Dd−/− NK cells and H2D^d^ negative cells in Dd+/+ and Dd+/− NK cells, and these cells were excluded for quality reasons **(Fig. S1d)**. Furthermore, one experiment contained too few Dd−/− cells and the analysis was hence based on two pooled datasets **(Fig. 1c-f)**. Dimensionality reduction by uniform Manifold Approximation and Projection (UMAP) on Principle Component Analysis (PCA) dimensions revealed a large overlap, as expected from a highly purified starting population **(Fig. 1c)**. Although no clear sub-cluster of cells from either mouse was observed, Dd−/− cells mostly localized to one end of the UMAP **(Fig. 1c, left panel)**.

Because of low experimental variability using the hash-tagging protocol, we initially set a low threshold to identify possible differences between the three samples. When comparing uneducated NK cells (Dd−/−) with educated NK cells (Dd+/− or Dd+/+), several genes were differentially expressed more than 1.2-fold, many of them previously implicated in key NK cell functions or maturation processes including *Id2, Klrg1, Ptpn6, Ccl5, Rap1b, Ltb*, and *Ctla2a* **(Fig. 1d, S1e, Supplementary table 2)** ^27, 28, 29, 30, 31, 32, 33, 34^. A large overlap in differently expressed genes (DEG) of the comparisons Dd−/− vs Dd+/+ and Dd−/− vs Dd+/− indicated that education by one MHC-I copy was sufficient to drive transcriptional changes **(Fig. 1d)**. Pathway analysis of DEG from the Dd−/− and Dd+/+ comparison revealed that all top pathways were immune response-related **(Fig. S1f)**. This confirmed our hypothesis that NK cell education is associated with discrete transcriptional changes.

Because the educational rheostat is controlled by different expression levels of H2D^d^, we next compared the transcriptional profiles of sp-Ly49A NK cells from Dd+/− and Dd+/+ mice. Differences between these two mice were limited to three transcripts: *Gm10076*, *Ub52* and *H2-T23* **(Fig. 1d)**. The two first were also overexpressed in Dd−/− mice when compared to Dd+/− mice but not when compared to Dd+/+ mice, suggesting an artefactual deficiency of these markers in our Dd+/− samples.

We next asked if the transcriptional differences associated with education were persistent across maturation by grouping the scRNA-seq data based on CD27 and CD11b expression ^35^. Differences between educated and uneducated NK cells were found in all three subsets (CD27^high^CD11b^low^, CD27^high^CD11b^high^ or CD27^low^CD11b^high^) with the largest number of DEGs in the least mature CD27^high^CD11b^low^ subset **(Fig. 1e)**. A few genes that were differentially expressed in the CD11b^high^ subset compared to the immature CD27^high^ subset were calcium binding proteins **(Fig. 1f, Supplementary table 3)**. Importantly, *Ptpn6* remained a commonly upregulated transcript in Dd−/− vs Dd+/+ NK cells in all subsets, indicating its independence from changes involved with maturation **(Fig. 1e, f)**. In summary, our data support the recently published findings of Wu and colleagues ^24^, who described increased *Ptpn6* transcription in uneducated NK cells. Furthermore, our analysis establish that altered *Ptpn6* transcription transgresses NK cell maturation.

### SHP-1 expression levels are linked to murine NK cell education

Quantitative PCR (qPCR) confirmed that *Ptpn6* transcripts were significantly more abundant in Dd−/− NK cells as compared to Dd+/− or Dd+/+ NK cells, but no difference was seen between Dd+/− and Dd+/+ mice **(Fig. 2a)**. All three strains showed similar transcript levels of *Ptpn11* and *Inpp5d*, which encode the two other main phosphatases in NK cells, SHP-2 and SHIP, and for two unrelated phosphatases, *Ptprs* and *Ppp4c* **(Fig. 2a)**. *Ptpn6* and *Ptpn11* were comparably expressed in Dd+/+ and Dd+/− NK cells relative to *Gapdh* **(Fig. S2a)**. Next, we used flow cytometry and a validated rabbit anti-mouse SHP-1 antibody to examine SHP-1 expression at the protein level in barcoded cells **(Fig. S2b and S2c)**. To dissociate NK cell education from the maturation process, we further stratified the data based on CD11b expression ^36^. Confirming our transcriptional analysis, Dd−/− NK cells expressed significantly higher levels of SHP-1 compared to Dd+/+ and Dd+/− NK cells **(Fig. 2b, left)**. We noted a small difference between Dd+/+ and Dd+/− NK cells, but it did not reach statistical significance. Interestingly, in all three strains, the less mature CD11b-NK cells expressed higher levels of SHP-1 as compared to the more mature CD11b+ subset **(Fig. 2b, right and Fig. 2c)**. Similar results were seen when including CD27 in the definition of maturation **(Fig. S2d)**. Expression of SHP-2 and SHIP were also significantly higher in immature as compared to mature NK cells, but in accordance with the qPCR data, no differences were seen between Dd−/− and Dd+/+ NK cells **(Fig. S2e, f)**. Further confirming the qPCR and scRNA-seq data, the difference between Dd+/− and Dd+/+ NK cells was not statistically different **(Fig. 2b, c)**. We conclude that the phosphatases SHP-1, SHP-2 and SHIP are all associated with NK cell maturation, yet only SHP-1 expression is regulated by H2D^d^-mediated NK cell education.

**Fig. 2.**
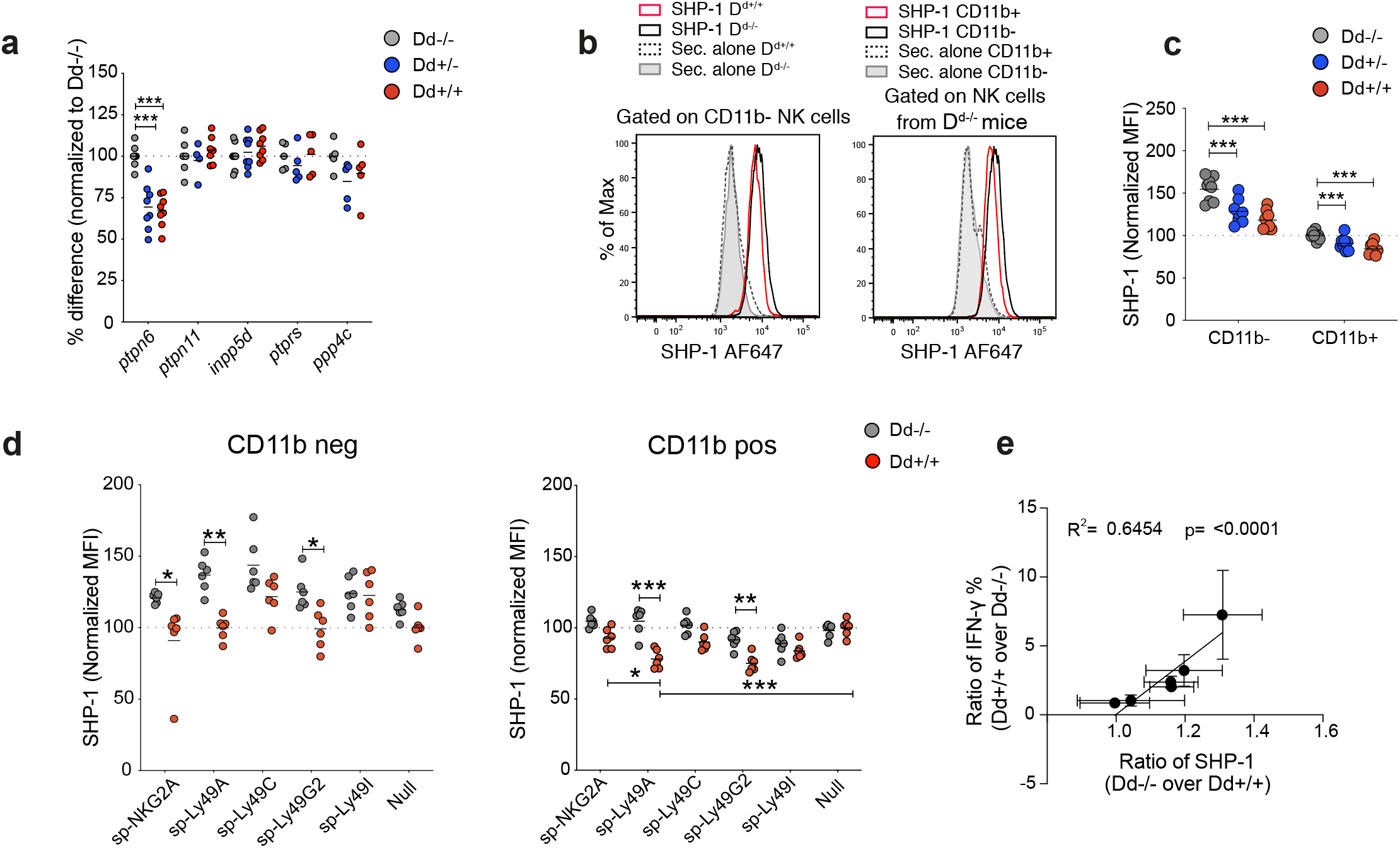
SHP-1 expression level is linked to murine NK cell education but not to the rheostat. **a,** Expression of phosphatases at mRNA level in NK cells. Data were pooled from four independent experiments (n = 5-8). Data were normalized to GAPDH and then to the average expression of Dd+/+ NK cells in each experiment. **b-c,** Representative (b) and summary (c) of SHP-1 expression at the protein level measured flow cytometry on CD11b- or CD11b+ NK cells. Data were pooled from three independent experiments (n=8). After subtracting the MFI of staining with secondary antibody alone, MFIs were normalized to the average MFI Dd−/− CD11b+ in each experiment. **d,** SHP-1 expression in NK cell subsets expressing one of the five main inhibitory receptors in immature (CD11b-, left panel) or mature (CD11b+, right panel) NK cells. Data were pooled from two independent experiments (n=6). Normalized MFI: MFI subtracted to secondary alone samples and normalized to the average of Dd+/+ null subset in each experiment. **e,** Correlation of differences in SHP-1 expression with differences in fractions of cells producing IFN-γ. The six subsets are those that express single inhibitory receptors or the null subsets from Dd−/− and Dd+/+ mice. Error bars: Means and SD. **a, c, d**, Two-way ANOVA with Sidak’s multiple comparisons. **e,** Linear regression. *p < 0.05, **p < 0.01, ***p < 0.001.

To further deepen the phenotypic analysis, we explored SHP-1 expression in CD11b- and CD11b+ NK cells expressing single inhibitory receptors **(Fig. S1b)**. In both CD11b- and CD11b+ NK cells, SHP-1 expression was lower in sp-NKG2A, sp-Ly49A, sp-Ly49G2 and sp-Ly49C NK cells from Dd+/+ compared to Dd−/− mice, while SHP-1 expression in sp-Ly49I and inhibitory receptor null NK cells was not affected **(Fig. 2d)**. Overall, the difference in SHP-1 expression between uneducated and educated NK cells for a given subset (expressed as a ratio) correlated well to the magnitude of education of the same subset, measured as a ratio between educated and uneducated NK cells with regards to the fraction of NK cell expressing IFN-g after stimulation **(Fig. 2e)**. In other words, the stronger the functional education by H2D^d^ on a given subset, the more pronounced was the reduction in SHP-1 expression in the same subset. We conclude that SHP-1 expression in NK cells is modulated both by NK cell maturation and MHC class I-binding inhibitory receptor interactions, resulting in an inverse correlation between function and SHP-1 protein expression.

### SHP-1 inhibition or gene deletion augments NK cell function in uneducated NK cells

The higher abundance of SHP-1 in uneducated NK cells might present a barrier downstream of activating receptors leading to a higher activation threshold and hypo-responsiveness. To test this hypothesis, we determined if treating Dd−/− NK cells with a chemical inhibitor of SHP-1 ^37^ would augment NK cell function. A significant increase in degranulation and IFN-g secretion was seen after inhibitor treatment compared to control treatment **(Fig. 3a)**. Next, we targeted SHP-1 in primary mouse NK cells using CRISPR-Cas9. Partial deletion of SHP-1 expression was seen at four days postelectroporation as compared to the control without SHP-1 crRNA **(Fig. 3b)**. Culturing NK cells for four days to some extent broke tolerance of Dd−/− NK cells, but despite this, SHP-1 reduction resulted in a higher NK cell response to NK1.1 stimulation **(Fig. 3c)**. Thus, inhibition or reduction of SHP-1 in uneducated Dd−/− NK cells potentiated activating receptor signaling, suggesting that SHP-1 expression contributes to hypo-responsiveness in uneducated NK cells.

**Fig. 3.**
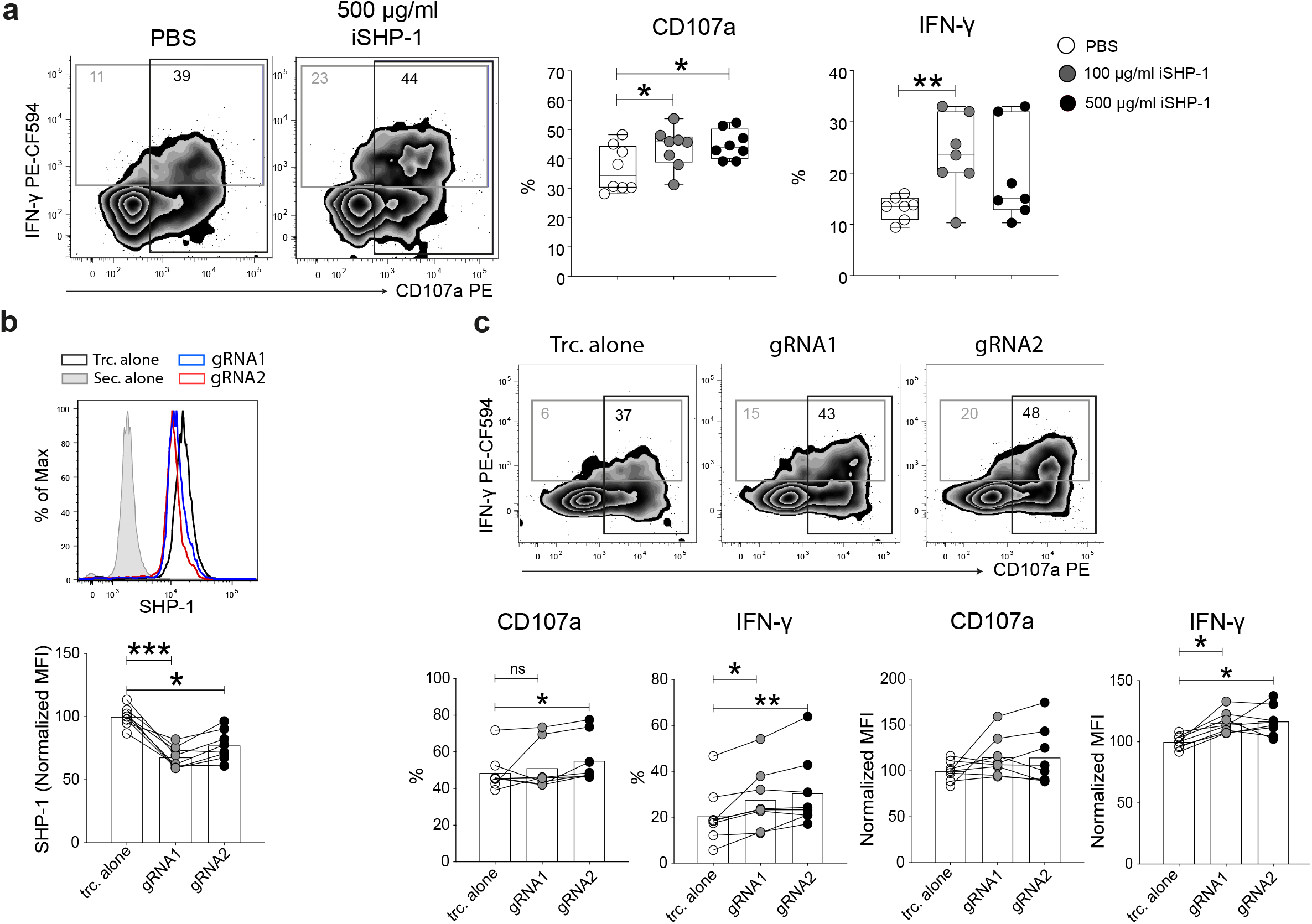
SHP-1 inhibition or knock-out augments uneducated NK cell function. **a,** NK cell degranulation and IFN-γ production upon NK1.1 stimulation after treatment with SHP-1 inhibitor (NSC-87877). Data were pooled from 3 independent experiments (n = 4-5). **b**, SHP-1 expression upon CRISPR/Cas9-mediated SHP-1 depletion. Enriched NK cells were activated for 4 days with 50 ng/ml rmIL-15 prior to electroporation with sgRNA and Cas9. Following electroporation, NK cells were cultured for another 4 days in the presence of 50 ng/ml rmIL-15 prior to functional assays. Data were pooled from five independent experiments (n = 8). **c**, IFN-γ production and degranulation triggered via NK1.1 stimulation upon SHP-1 KO. Data were pooled from four independent experiments (n =8). Error bars: Means and SD. One-way ANOVA with Dunn’s multiple comparisons. *p < 0.05, **p < 0.01.

### SHP-1 expression in human NK cells is defined by the education status

Differences between NK cell education in humans and mice have been reported ^26, 38, 39, 40^ and it was therefore of interest to investigate if SHP-1 was expressed to higher levels in uneducated NK cells also in humans. The specificity of a monoclonal rat-anti human SHP-1 antibody was confirmed by CRISPR-Cas9 mediated *PTPN6* KO **(Fig. S3a)**. Functional assays with anti-CD16 crosslinking confirmed that educated subsets (self-KIR, NKG2A+) produced more IFN-γ and degranulated more than uneducated subsets (nonself-KIR, NKG2A-KIR-) **(Fig. S3b and Fig. 4a)**. Corroborating the mouse data, uneducated NK cells (non-self KIR+ or NKG2A-KIR-) expressed higher SHP-1 levels as compared to educated subsets (NKG2A+ or self-KIR+) **(Fig. 4b, c)**. Further, iSHP-1 treatment augmented the capacity of human NK cells to produce IFN-γ and degranulate in response to CD16 stimulation **(Fig. 4d)**. We conclude that SHP-1 was more highly expressed in uneducated compared to educated human NK cells and was a barrier to activation of effector functions.

**Fig. 4.**
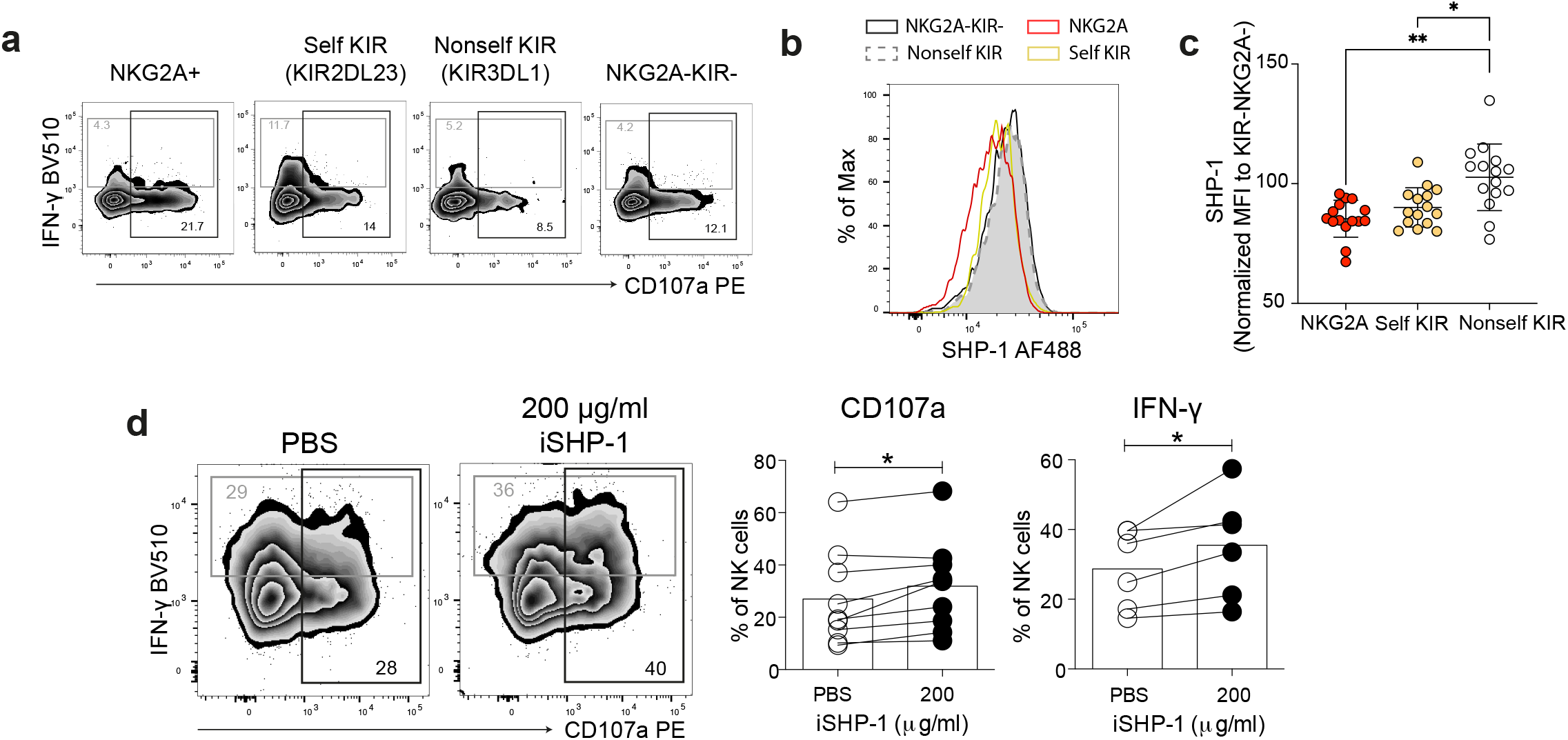
SHP-1 expression is associated with human NK cell education and modulation of SHP-1 activity or expression changes NK cell functions. **a**, Functions of NKG2A-KIR-, nonself KIR3DL1+, self KIR2DL23+ and NKG2A+ NK cells upon anti-CD16 plate-bound stimulation. **b, c,** Representative plot (b) and summary (c) showing SHP-1 expression in the subsets mentioned in **a**. Data were pooled from 6 independent experiments (n = 19). **d**, Effects of iSHP-1 (NSC-87877) on IFN-γ production and degranulation of NK cells triggered via CD16 stimulation. Data were pooled from five independent experiments (n = 6-9). **c**, One-way ANOVA with Dunn’s multiple comparisons. **d**, Mann-Whitney tests. *p < 0.05, **p < 0.01.

### NK cell education reduces constitutive SHP-1 activity and enhances NK cell signaling

During NK cell inhibition mediated by MHC class I-specific receptors, recruited SHP-1 becomes activated and triggers dephosphorylation of several nearby signaling molecules such as VAV-1, LAT, SLP76, ZAP70 and PLC-g2, limiting NK cell responses ^41, 42, 43, 44^. We hypothesized that an abundance of SHP-1 in uneducated NK cells might not require inhibitory receptor crosslinking to limit signaling downstream of activating receptors compared to educated NK cells. To test this, we stimulated educated and uneducated NK cells with an antibody again NKp46 and performed phosphoflow analysis. A cellular barcoding protocol was again used to minimize experimental variability **(Fig. S4a)** ^45^. Besides signaling molecules implicated as direct SHP-1 targets, we also examined activation of ERK1/2, a down-stream read-out of NK cell activation ^46^.

Phosphorylation of ERK1/2 (pT202/pY204) and PLC-γ2 (pY759) was significantly increased upon NKp46 stimulation of Ly49A+ NK cells from Dd+/+ mice but not from Dd−/− mice **(Fig. 5a, b)**. Phosphorylation of SLP76 (pY128) and ZAP70/SYK (pY319/pY352) was induced by NKp46 stimulation in Ly49A+ NK cells from both Dd−/− and Dd+/+ mice, yet with higher increment ratios in Dd+/+ NK cells **(Fig. 5c and S4b)**. As expected, the activation of STAT5 at Y694 was not induced by NKp46 stimulation irrespective of mouse strain, serving as a negative control **(Fig. 5d)**. To test if the lower activation of SLP76, PLC-g2, ZAP70/SYK and ERK1/2 was linked to SHP-1 activity in Dd−/− NK cells, we performed the same experiment in the presence of iSHP-1. Inhibitor-treated Dd−/− NK cells indeed showed augmented phosphorylation of all four signaling molecules to different extents, providing direct evidence for a dampening role of SHP-1 on NK cell signaling in uneducated NK cells **(Fig. 5e)**. No similar increase was seen in Dd+/+ NK cells **(Fig. 5e)**, implying qualitative differences in the way SHP-1 may control activating receptor signaling in educated and uneducated NK cells.

**Fig. 5.**
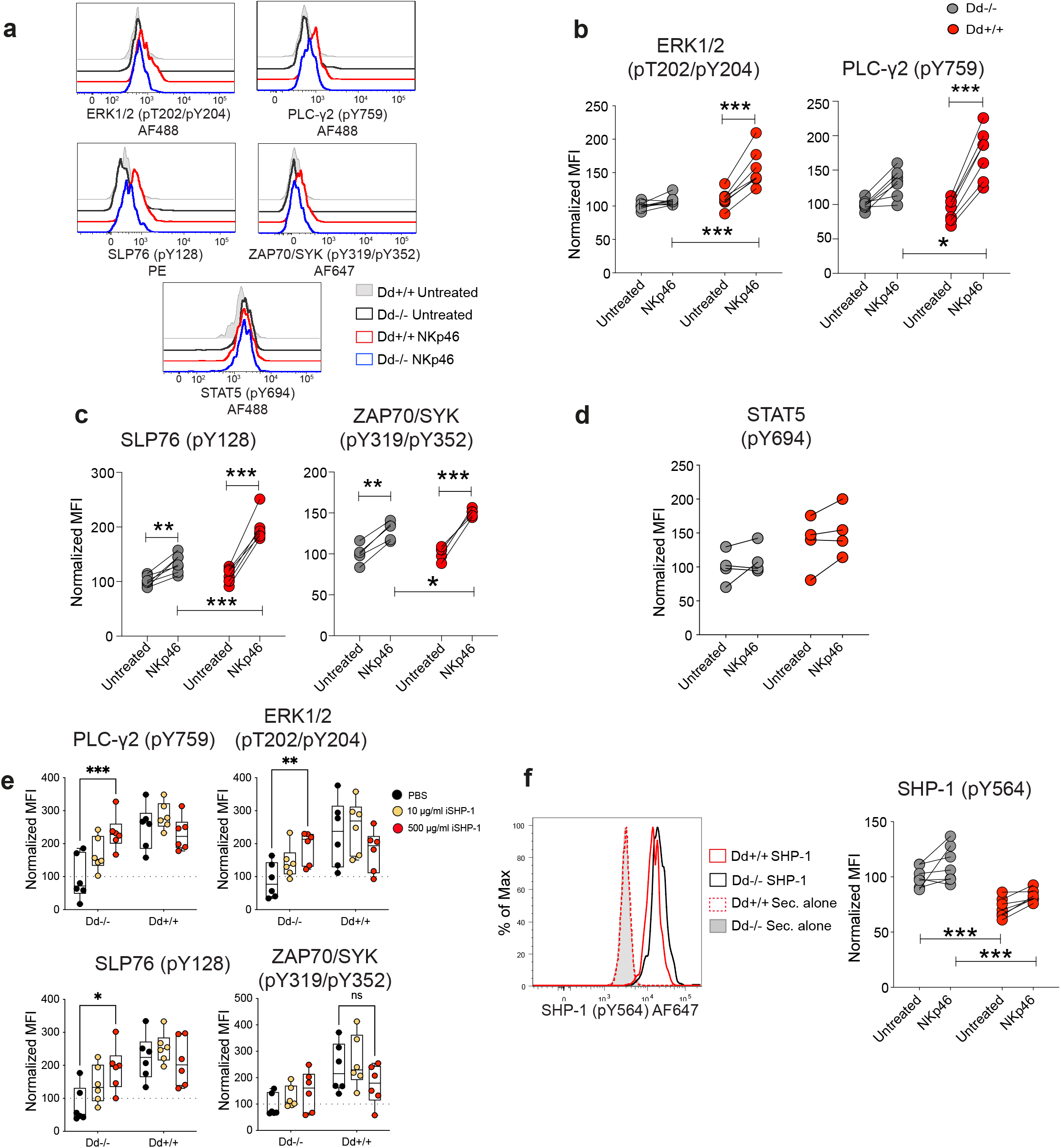
Uneducated NK cells possess higher level of activated SHP-1 and lower phosphorylation status of SHP-1 substrates. **a-d**, Representative flow cytometry histograms **(a)** and summary data **(b-d)** of ERK1/2 (pT202/pY204), PLC-γ2 (pY759), SLP76 (pY128), ZAP70/SYK (pY319/pY352), STAT5 (pY694) in steady state and after NKp46 stimulation. **e,** Phosphorylation of these signaling molecules upon iSHP-1 treatment and NKp46 stimulation. Normalized MFI: After subtraction of the MFI of isotype control staining, MFIs were normalized to the average of PBS treated Dd−/− NK cells in each experiment. Data were pooled from 3 independent experiments (n = 6). **f**, Active SHP-1 (pY564) in steady state and upon NKp46 stimulation. **b-d**,**f**, Normalized MFI: After subtraction of the MFI of isotype control staining, MFIs were normalized to the average of untreated Dd−/− NK cells in each experiment. Data were pooled from 2-3 independent experiments (n = 4-7). Error bars: Means and SD. **b-f**, Two-way ANOVA with Sidak’s multiple comparisons, ns: non-significant, *p < 0.05, **p < 0.01, ***p < 0.001.

The phosphatase activity of SHP-1 is itself regulated by phosphorylation. The N-terminal SH2 domain of SHP-1 serves as an intramolecular switch, binding to the phosphatase domain and inhibiting its activity ^47, 48^. However, when the SHP-1 SH2 domain binds to phosphorylated ligands, including intramolecularly to C-terminal (Y536 and Y564) SHP-1 phosphorylated sites, the phosphatase domain is released from inhibition, thus becoming active ^47, 48^. Phosphorylation of the SHP-1 Y536 and Y564 residues have been associated with an increased phosphatase activity ^49^. Remarkably, phosphoflow experiments demonstrated that the phosphorylation of SHP-1 (pY564) was significantly higher in Dd−/− NK cells at steady state compared to NK cells from Dd+/+ mice **(Fig. 5f)**. NKp46 stimulation increased, yet not significantly, phosphorylated SHP-1 (pY564) levels in both Dd−/− and Dd+/+ NK cells **(Fig. 5f)** but the main difference was seen at steady state. Altogether, these data suggest that a higher amount of constitutively activated SHP-1 in uneducated NK cells is associated with reduced activating receptor signaling following AR stimulation.

### Educated NK cells show less SHP-1 and more phosphorylated SLP76 (pY128) in the activating synapse

Previous studies have implied that SHP-1 localization in the immune synapse may be dynamically controlled ^50, 51, 52^ and we therefore asked if SHP-1 distribution in the NK cell activating synapse was different in educated and uneducated NK cells **(Fig. S5a)**. An activating synapse was induced by incubating enriched NK cells on anti-NKp46 antibody-coated glass coverslips, followed by confocal imaging of the interaction site formed between the NK cells and the antibody-coated glass. Ly49A+ NK cells from Dd−/− mice formed smaller synaptic interfaces **(Fig. 6a,b)** and were less compact as compared to Dd+/+ Ly49A+ NK cells **(Fig. 6c),** consistent with a non-activated state.

**Fig. 6.**
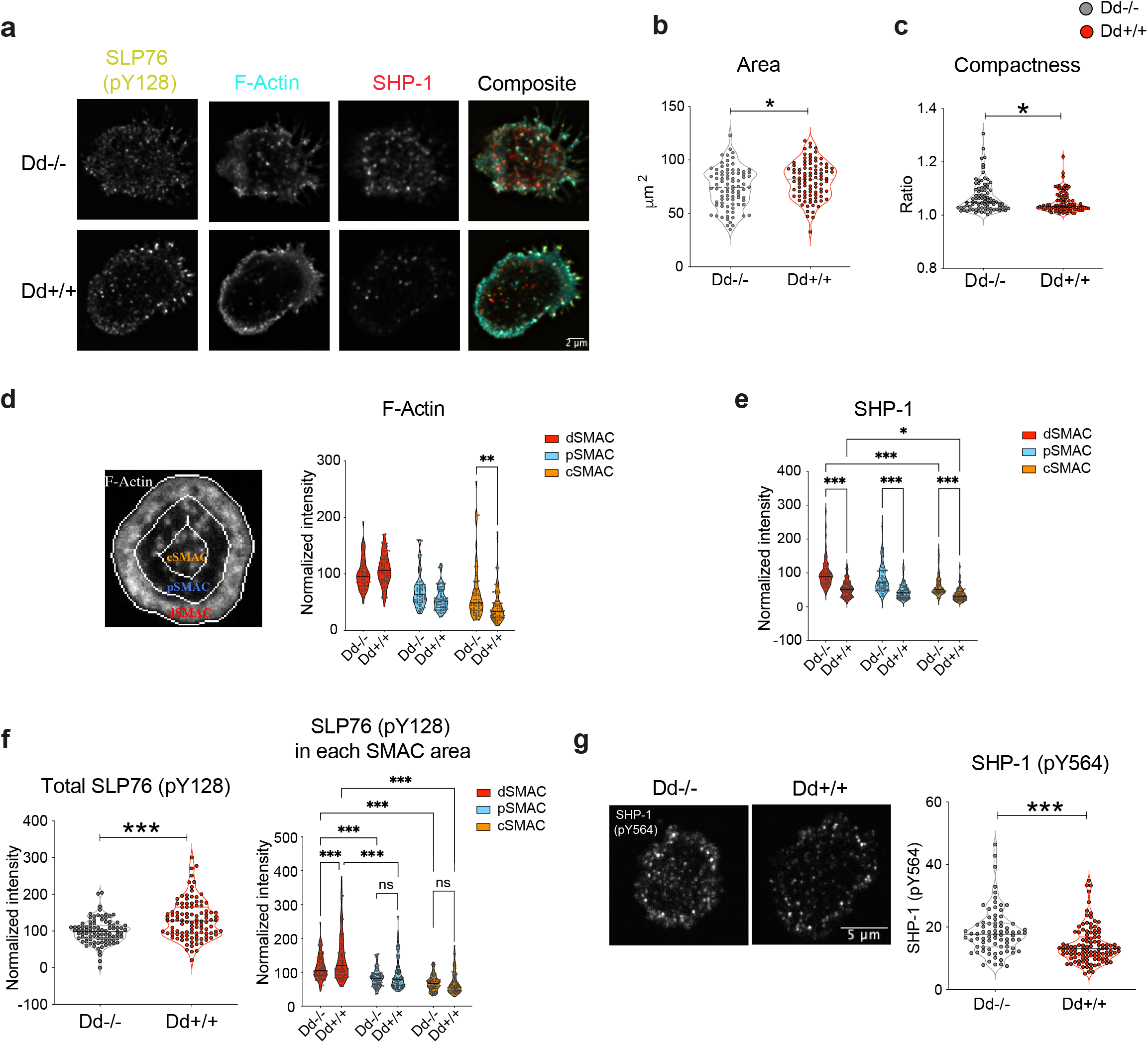
SHP-1 and SLP76 (pY128) localization in the activating synapse of NK cells from Dd−/− and Dd+/+ mice. **a**, Representative images of phospho-SLP76 (pY128), SHP-1 and F-actin on NKp46 synapse of Ly49A+ NK cells. **b-c**, Quantification of the area of the synapses (**b**) and the synapse compactness (**c**) of cells from **a.** Data were pooled from two independent experiments (n=60-67). **d-e**, Representative image showing the synaptic regions defined for image analysis (**d** left), quantification of the phalloidin (F-actin) fluorescence intensity for each synaptic region (**d** right). SHP-1 intensity in cSMAC, pSMAC and dSMAC is showed in (**e**) Data were pooled from three independent experiments (n=90-104). **f**, SHP-1 (pY564) intensity on NKp46 synapse of sp-Ly49A cells. Data were pooled from two independent experiments (n=95-111). **h**, Phosopho-SLP76 (pY128) on NKp46 synapse of Ly49A+ cells. Data were pooled from three independent experiments (n=90-104). Error bars: Means and SD. **d, e, f right panel,** Two-way ANOVA with Sidak’s multiple comparisons post-tests. **b**, **c**, **g right panel** Mann-Whitney tests. ns: non-significant, *p < 0.05, **p < 0.01, ***p < 0.001.

Activating immunological synapses formed by cytotoxic T cells and NK cells are characterized by organization of the cell surface into supramolecular activation clusters (SMACs) and concomitant reorganization of the underlying cortical actin cytoskeleton that serves to regulate activation and facilitate synaptic processes ^53^. To gain an understanding of how SHP-1 activity might regulate NK cell activation, we identified the different areas of the synaptic interface, i.e. distal (dSMAC), peripheral SMAC (pSMAC) and central SMAC (cSMAC), based on the F-actin organization. Specifically, the outermost dense actin-ring (dSMAC), the less dense lamellar actin residing inside the dSMAC (pSMAC) and the innermost actin hypodense area (cSMAC) were studied. Consistent with visual inspection of images, analysis of actin enrichment in each area revealed that the synapses formed could be characterized by the actin enrichment in dSMAC **(Fig. 6d)**. In line with the modest increase in cellular spreading, Dd+/+ NK cells accumulated more dSMAC F-actin. Conversely, in uneducated Dd−/− NK cells, there was a less pronounced differential enrichment in the SMACs with significantly more F actin observed in the cSMAC **(Fig. 6d)**.

Analysis of SHP-1 and phospho-SLP76 (pY128) in the different SMACs revealed that both molecules were enriched in the actin-rich dSMAC followed by pSMAC regardless of the mouse strains **(Fig. 6e,f)**. In the dSMAC area, SHP-1 staining was denser while phospho-SLP76 (pY128) staining was weaker in Dd−/− vs Dd+/+ synapses, indicating their negative correlation **(Fig. 6e,f)**. To determine whether SHP-1 that was recruited to Dd−/− and Dd+/+ activating NK cell synapses was functionally active, we examined the presence of active, phosphorylated SHP-1. In both cases, active SHP-1 (pY564) was recruited to the synaptic interface with a significantly higher density at sp-Ly49A Dd−/− NK cell synapses **(Fig. 6g)**, which is in agreement with the phosphoflow data **(Fig. 5e)**. Our data suggest that the activating synapse of the uneducated NK cells bears resemblance to inhibitory synapses, in which SHP-1 is recruited by inhibitory receptors ^51^. These similarities lend further support to a model in which the recruitment of SHP-1 to the synapse of uneducated NK cells is a key step in dampening signaling, thereby maintaining self-tolerance.

### SHP-1 forms close contacts with actin and SLP-76 in the activating synapse of uneducated NK cells

In the T cell synapse, F-actin foci colocalize with T cell receptor (TCR) micro-clusters crucial for activating signaling and calcium mobilization ^54^. We hypothesized that, in NK cells, SHP-1 may be localized in proximity to similar activating clusters and to actin, thereby attenuating activating signal transduction. To test this hypothesis, we developed a super-resolution imaging protocol using stimulated emission depletion (STED) microscopy to gain insight in the localization of SHP-1, SLP76 (pY128) and F-actin in the synapse. In NKp46-activating synapses of Dd−/− NK cells as compared to Dd+/+ NK cells, a higher degree of overlap between SHP-1 and phospho-SLP76 (pY128) or SHP-1 and F-actin was observed, as measured by image-based colocalization levels (Manders coefficient) **(Fig. 7a,b)**. We next measured the actual distance between SHP-1 and phospho-SLP-76 (pY128) clusters, by transforming SHP-1 objects into distance maps and calculating the minimum distances of phospho-SLP-76 (pY128) to surrounding SHP-1. This distance was significantly lower in the Dd−/− synapse compared to the Dd+/+ synapse **(Fig. 7c)**, with most objects showing a complete overlap with a distance around 0 μm. Some far away clusters were also observed that were more enriched in Dd+/+ synapses **(Fig. 7c)**. Similarly, STED images of phospho-SHP-1 (pY564) revealed a close association of activated SHP-1 with F-actin-enriched regions and SLP76 **(Fig. 7d, e)**, with a similar high degree of direct overlap between phospho-SHP-1 (pY564) and F-actin or SLP76 **(Fig. 7d, e),** as when the entire SHP-1 protein was imaged **(Fig. 7c)**. A similar object-based distance analysis revealed a closer distance between SLP76 to surrounding phospho-SHP-1 (pY564) objects **(Fig. 7f).** Taken together, our data reveal that SHP-1 and activated SHP-1 reside in proximity to SLP76 and phosphorylated SLP76 (pY128) in actin-enriched regions in the activating synapse of uneducated NK cells, providing a plausible explanation for the increased dampening effect of AR signaling in uneducated NK cells that contain higher abundance of SHP-1.

**Fig. 7.**
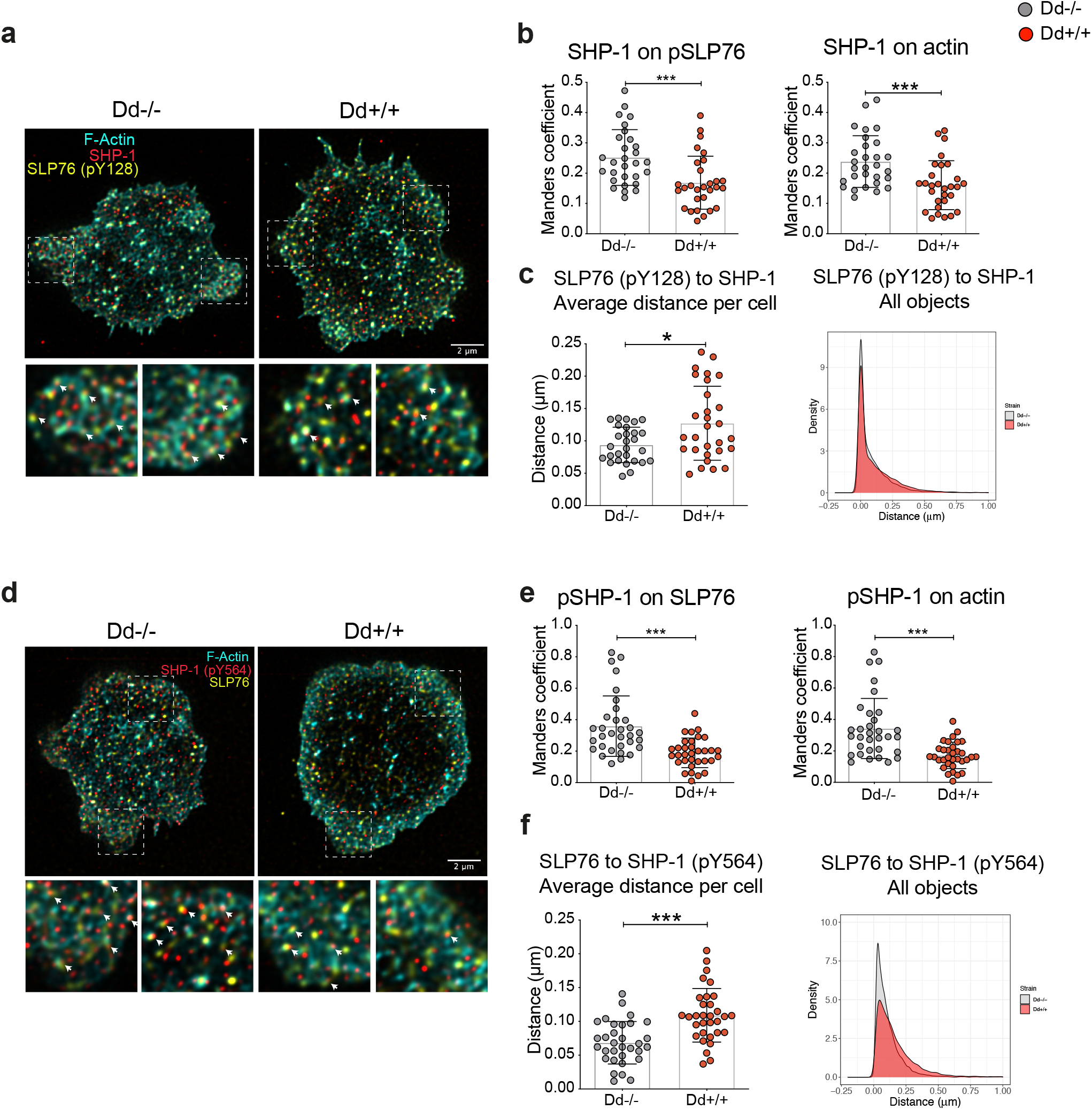
STED microscopy demonstrates a higher enrichment of SHP-1 next to F-actin rich regions and SLP76 (pY128) clusters on the activating synapse of Dd−/− NK cells. **a**, Representative STED images of SHP-1, F-actin and phospho-SLP76 (pY128) on NKp46 synapse. **b,** Image-based colocalization analysis of SHP-1 and F-actin or SHP-1 and phospho-SLP76 (pY128) from **a**. **c,** Object-based distance analysis showing the minimum distance of phospho-SLP76 (pY128) objects to surrounding SHP-1 from **a**. Average distance per cell (left panel) and density plot showing distance of all objects (right panel). Data was from one out of two independent experiments (n=29-31). **d,** Representative STED images of phospho-SHP-1 (pY564), F-actin and SLP76 on NKp46 synapse. **e,** Image-based colocalization analysis of SHP-1 and F-actin or SHP-1 and phospho-SLP76 (pY128) from **d**. **f,** Object-based distance analysis showing the minimum distance of SLP76 objects to surrounding phospho-SHP-1 (pY564) from **d**. Average distance per cell (left panel) and density plot showing distance of all objects (right panel). Data was from one out of two independent experiments (n=32-33). Error bars: Means and SD. **b, c, e, f**, Mann-Whitney tests, *p < 0.05, ***p < 0.001.

### Uneducated Ly49A+ NK cells display low association between SHP-1 and Ly49A in the cell membrane and enhanced enrichment of SHP-1 in the activating synapse

Increased localization of SHP-1 in the activating synapse of uneducated NK cells could be an indirect consequence of higher SHP-1 abundance overall or a direct mechanism promoting localization of SHP-1 to the activating synapse at the expense of the rest of the cell. To investigate the latter possibility, we quantified SHP-1 enrichment in the NKp46-activated synapse by measuring the ratio of SHP-1 in the 2 μm-thick synapse relative to that in the whole cell. This analysis revealed a significantly higher synapse enrichment ratio of SHP-1 for Dd−/− NK cells **(Fig. 8a,b)**, supporting the existence of an active mechanism in the uneducated NK cell synapse upon activation. Interestingly, this finding was paralleled with an enrichment **(Fig. 8a,c)** and increased absolute amount **(Fig. 8c,d)** of Ly49A in the uneducated synapse, suggesting an involvement of Ly49A in the SHP-1 enrichment process. A possible explanation for reduced SHP-1 enrichment in the activating synapse of educated NK cells might be SHP-1 sequestration, of which the educating inhibitory receptor in Dd+/+ NK cells Ly49A is a possible candidate ^55^. To determine if this was the case, we performed STED microscopy with resting cells cyto-spinned onto coverslips. Indeed, SHP-1 was significantly more colocalized with Ly49A in Dd+/+ sp-Ly49A NK cells as compared to those from Dd−/− mice **(Fig. 8e)**, inferring that educating inhibitory receptors might sequester SHP-1, contributing to a lower SHP-1 level at the activating synapse.

**Fig. 8.**
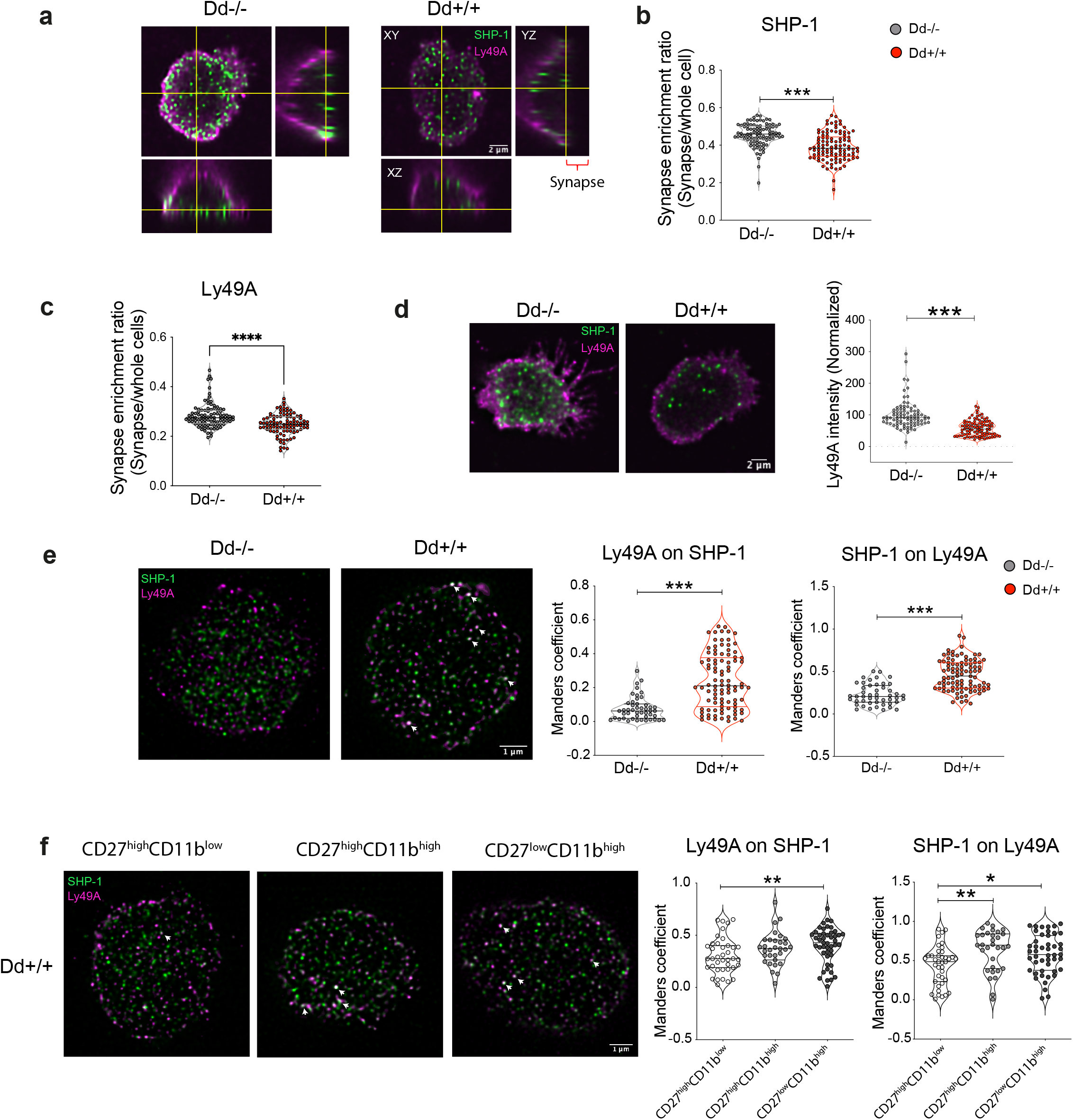
SHP-1 sequestration by educating inhibitory receptor on Dd+/+ NK cells. **a,** Representative images of SHP-1 and Ly49A distribution over NKp46 activated cells. **b, c**, Enrichment ratios of **(b)** SHP-1 and **(c)** Ly49A intensity in the synapse area over the whole cells. Data were pooled from two independent experiments (n=95-111). **d,** Ly49A intensity in the NKp46 synapse. **e,** STED images SHP-1 and Ly49A on the cell membrane of resting sp-Ly49A NK cells from Dd−/− and Dd+/+ mice. Data were pooled from four independent experiments (n =47-94). Image-based colocalization analysis of SHP-1 and Ly49A on the cell membrane of resting sp-Ly49A NK cells stratified based on different maturation stages from Dd+/+ mice. The data was from one experiment (n = 32-47 in each group). Error bars: Means and SD. **b-e,** Mann-Whitney tests. **f,** Kruskal-Wallis with Dunn’s multiple comparisons. *p < 0.05, **p < 0.01, ***p < 0.001.

Finally, we asked whether the association between Ly49A and SHP-1 was affected by NK cell maturation. We sorted resting sp-Ly49A NK cells belonging to three maturation subsets, namely CD27^high^CD11b^low^, CD27^high^CD11b^high^ and CD27^low^CD11b^high^ **(Fig. S6)**, and performed STED microscopy. SHP-1 and Ly49A colocalization in Dd+/+ sp-Ly49A NK cells was increased along the maturation process **(Fig. 8c)**, following NK cell functional capacity. Together, these findings suggest that educating inhibitory receptors sequester SHP-1, contributing to a lower SHP-1 level at the activating synapse that directly affects NK cell function.

## DISCUSSION

Upon NK cell activation, an immune synapse is formed at the interphase between the NK cell and the target cell, facilitating sustained signaling, polarization of cytotoxic granules and release of their content to the synaptic cleft ^56^. Compared to the activating synapse, the inhibitory NK cell synapse is less organized, smaller, lacks a dense ring of F-actin in the p-SMAC and possesses a higher density c-SMAC actin mesh with smaller holes that are not penetrable for cytotoxic granules ^57, 58, 59, 60^. It has been posited that IR binding to facilitates recruitment of phosphatases such as SHP-1 to counteract early activation signals such as phosphorylated VAV-1 leading to actin reorganization ^41^. Following such inhibition, activation of other downstream signaling may be blocked, some of which may also represent direct targets of SHP-1 including Lck ^61^, ZAP70 ^44^, SLP-76 ^43^, PLC-γ ^42^ and LAT ^42^. Our experiments suggest that NK cell tolerance is controlled by SHP-1-mediated dephosphorylation of signaling molecules in uneducated NK cells, though augmented recruitment of SHP-1 to the activating immune synapse. The close association between SHP-1, actin and signaling molecules that we uncover in the uneducated immune synapse highlights similarities between the uneducated and the inhibitory NK cell synapse. We thus propose an integrated view of NK cell tolerance and education, positing a new model for how SHP-1 quantitatively controls NK cell education **(Fig. 9)** ^20, 24, 34^.

**Fig. 9.**
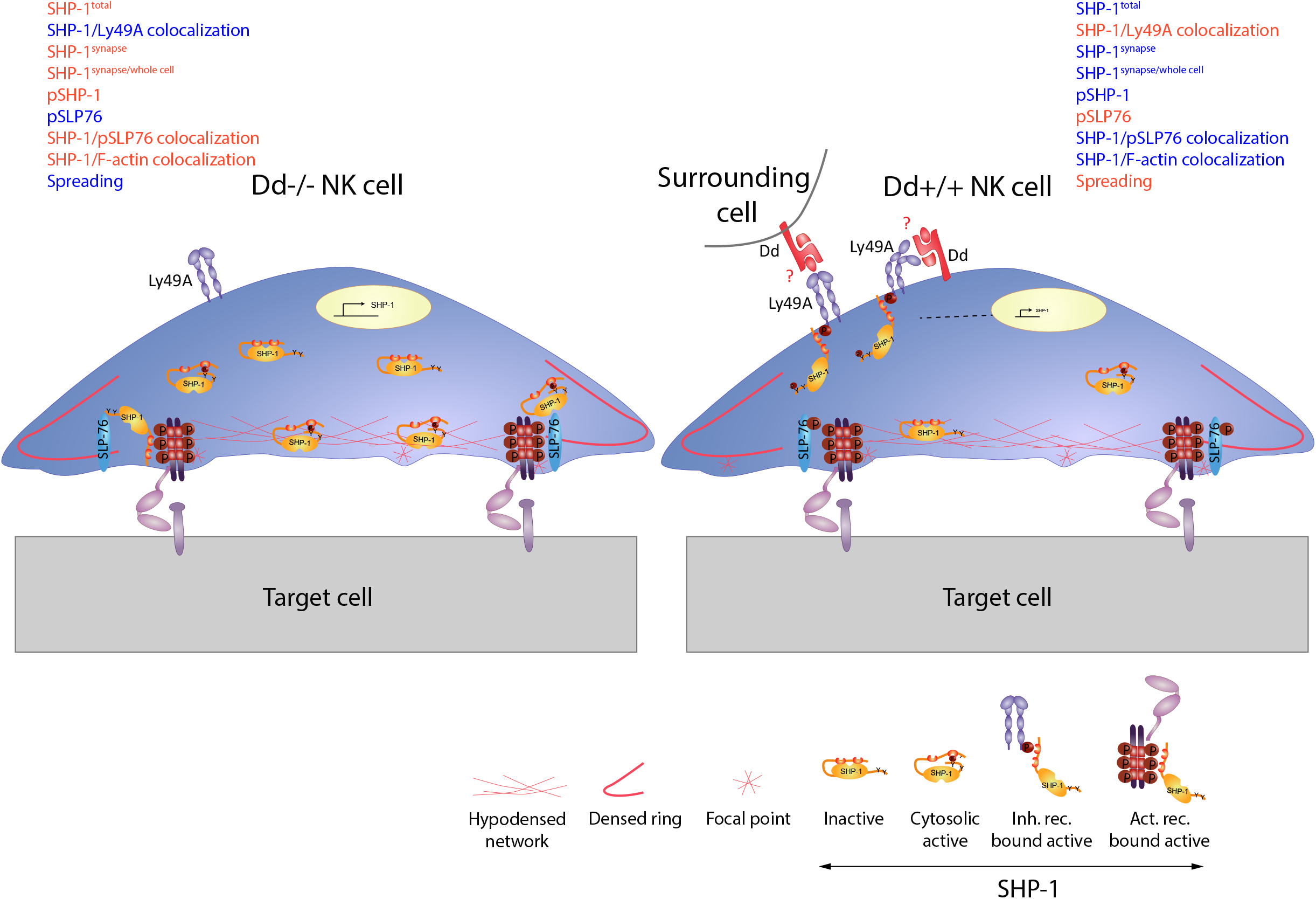
Model of SHP-1 as an important factor contributing to functional differences between educated and uneducated NK cells. The interaction between educating inhibitory receptor (Ly49A) with Dd lowered SHP-1 transcription in an unknown process. Ly49A/Dd interaction, wither in *cis* or in *trans*, recruits SHP-1, contributing to sequestering SHP-1 from the activating synapse. In addition, active SHP-1 (pY564) is higher in the Dd−/− synapse. SHP-1 is localized more closely with SLP76 (pY128) in Dd−/− NK cells. Consequently, SLP76 (pY128) is lower in Dd−/− synapse. In addition, SHP-1 is localized to a higher degree with F-actin-rich regions in Dd−/− cells, which might modulate SHP-1 conformation and activity. Taken together, as compared to Dd+/+ NK cells, those from Dd−/− mice are less or slower activated and respond weaker to activating receptor stimulation. Red letters: reduced levels, Blue letters: increased levels.

How is SHP-1 recruited to the activating synapse of uneducated NK cells in the absence of IR engagement? In our model we propose that the Ly49A receptor recruits SHP-1 to the membrane and restricts its synapse localization. We base this proposition on the skewed ratio between SHP-1 localization in the synapse and in the distal pole in uneducated NK cells and the parallel finding that Ly49A and SHP-1 were strongly colocalized in the cell membrane of educated but not uneducated NK cells. Whether or not the educating ligand H2D^d^ triggers the association between Ly49A and SHP-1 in our model is not known, but we deem this likely as education per definition requires an interaction between inhibitory receptors and their MHC class I ligands ^12, 29, 60^ Interactions between inhibitory receptors and their MHC class I ligands primarily take place in *trans*, between cells, but can also occur in *cis*, in the NK cell’s own membrane ^62^. A *cis* interaction has been described for many Ly49 receptors, including Ly49A ^63, 64^ and induces a conformation change in the Ly49A receptor ^65^. It thus represents an attractive aspect of our model for NK cell education since it would, at least theoretically, have a potential to restrict movement of the inhibitory receptor, with its associated SHP-1 phosphatase, to the synapse. In fact, a mechanism for NK cell tolerance involving a *cis* interaction between Ly49A and H2D^d^ has already been suggested by Werner Held and his group, who concluded that the *cis* interaction between Ly49A and H2D^d^ in educated NK cells restricts the localization of Ly49A to the interphase between NK cells and target cells ^55, 66^, data that were recapitulated by our finding in this paper of Ly49A being strongly enriched in the synapse of Dd−/− Ly49A+ NK cells. Whether or not SHP-1 is transported to the activating synapse by Ly49A in uneducated NK cells remains to be investigated. An equally likely model is that free SHP-1 enters the synapse of uneducated NK cells by itself upon NK cell activation, regulated in that case by a combination of increased abundance ^24^ and lack of Ly49A sequestration.

Not all inhibitory receptors are thought to bind their MHC class I ligands in *cis*, including some Ly49 receptors, NKG2A and KIR. However, this notion was challenged a few years ago for KIR3DL1 using a humanized mouse model, demonstrating binding to HLA-B*27:05 (containing the Bw4 epitope recognized by this KIR receptors) in *cis* ^25^. Importantly, the model we propose does not restrict itself to a potential *cis* interaction. If *cis* interactions do not take place, other mechanisms may operate in which an educating signal in *trans* leads to SHP-1 binding and sequestration outside of the synapse. Future work will be set up to test this possibility.

When Viant et al. deleted SHP-1 specifically in NK cells (Ncr1^iCre^ SHP-1^flox/flox^ mice), a markedly hypo-responsive NK cell phenotype developed that was very similar to that of β_2_m-deficient and MHC-I-deficient NK cells ^29^. This result is difficult to interpret in the light of our model defining SHP-1 as the foremost gatekeeper of tolerance in the uneducated immune synapse. One possibility to reconcile our model for tolerance with this data is to postulate that other phosphatases, for example SHP-2 and SHIP-1, can take the place of SHP-1 in the synapse, mediating similar dephosphorylation of activating kinases. Another possibility is that activating signals in SHP-1-deficient NK cells are attenuated by other mechanisms than phosphatase activity, for example by degradation of signaling adaptors^67^ or by exhaustion of the activating machinery and depletion of cytotoxic granule content, such as granzyme B, via unopposed signaling^68^.

In contrast to complete deletion, several studies have investigated how modulations or mutations of SHP-1 affect NK cell function and education. NK cells from SHP-1 defective motheaten-viable mice showed slightly lower IFN-γ production after NK1.1 stimulation ^8^. In another study, NK cells from motheaten-viable mice and mice expressing a catalytically inactive mutant of SHP-1 were hyporesponsive towards MHC class I-deficient cells but remained functional to antibody-dependent cell-mediated cytotoxicity^69^. More recently, SHP-1 knock down (KD) experiments *in vitro* yielded opposite data. ShRNA-mediated SHP-1 KD of primary mouse NK cells resulted in defective inhibitory signaling and an increased spontaneous degranulation and self-killing^70^. Furthermore, CRISPR/Cas9 mediated SHP-1-depletion or expression of inactive SHP-1 mutant in the YTS human NK cell line lowered inhibitory receptor signaling and increased HLA-positive tumor control *in vivo*^71^. In line with this data, we showed that CRISPR/Cas9 mediated SHP-1 depletion augmented the functional responsiveness of uneducated NK cells. Altogether, these data support a role for SHP-1 in controlling activating signaling in NK cells, consistent with our model.

What remains to be explained is how relative changes in the amount of SHP-1 protein could play dramatic and quantitative effects on NK cell responsiveness? In our hands, SHP-1 expression in educated NK cells is reduced by approximately 40% compared to uneducated NK cells, which means that also in educated NK cells, a large pool of SHP-1 is à priori available for possible recruitment to the activating synapse. Apparently, this does not happen as efficiently, suggesting that one or several active mechanisms, some of which we suggest here, may act in concert with transcriptional regulation in controlling the rheostat.

An intriguing question is how the mRNA/protein abundance is regulated by education? Wu et al. could not identify changes in chromatic accessibility of the SHP-1 gene conveyed by education^24^, leaving the mode of transcriptional regulation unsolved. An alternative way to explain the control of RNA/protein abundance by education may be to postulate a regulated RNA degradation process, quantitatively regulated by the strength of the educating signal according to the rheostat model. Such a mechanisms, if it exists, would be in direct equilibrium with the input educating signal and would lead to the conclusion that *Ptpn6* RNA abundance is a by-product of education rather than the reason for it. Also arguing against a direct causative role of the RNA abundance in our system is our finding that Dd+/− mice had the same RNA and very similar protein content (using both RNA sequencing, qPCR and flow cytometry) as Dd+/+ mice, despite the comparison of these two mouse models represent the essence of rheostat control of NK cell function. We also noted that NK cell maturation was associated with changes in SHP-1 expression independently from education. Thus, the more immature CD11b-NK cell subset expressed much more SHP-1 compared to mature NK cells. An interesting speculation is that SHP-1 is also involved in controlling the functional capacity of NK cells during maturation, with immature NK cells being less functional compared to mature NK cells. This question can be tested using imaging of activated SHP-1 in the immune synapse of mature and immature NK cells.

Irrespective of the mechanism for SHP-1 enrichment in the uneducated synapse, a speculative possibility is that it becomes activated by binding to actin, which under the slow retrograde flow characterizing the actomyosin dynamics might bring SHP-1 to the right place and offer binding sites for SHP-1 leading to SHP-1 phosphorylation and functional consequences for signaling ^50, 72^. Uneducated NK cells indeed contained a larger pool of active SHP-1 (pY564) compared to educated NK cells, and using STED microscopy, we identified high amount of SHP-1 (pY128) in the synapse in micro-clusters including activating receptors, activating signaling molecules and actin. Thus, not only do educated NK cell have less SHP-1 compared to uneducated NK cells, their pool is also less active. The close association of SHP-1 with F-actin and activating clusters as well as its enrichment in the d-SMAC might thereby provide a mechanism by which activating signals are dampened when activating receptors and signaling clusters migrate to the inside of the synapse from the d-SMAC ^73^.

In conclusion, we propose that NK cell tolerance is regulated by a mechanism similar to that mediating NK cell inhibition following MHC class I engagement. We propose that SHP-1 recruitment to the synapse following NK cell activation disarms activating signals and secures NK cell tolerance. In our model **(Fig 9)**, we propose that recruitment of SHP-1 to the synapse is restricted by SHP-1 sequestration outside of the synapse by the Ly49A receptor, following binding to its MHC-I ligand H2D^d^. Different expression levels, or different binding affinities to MHC-I, quantitatively control NK cell education. If SHP-1 sequestration is linearly regulated by MHC-I engagement, our model would provide an explanation for quantitative NK cell education. This and several other open questions are next in line to be addressed. Finally, we note that our model for tolerance is different from previously suggested models of general NK cells exhaustion from continuous encounters by activating receptor ligands ^68, 74, 75, 76^. Hypo-responsiveness based on recruitment of active phosphatases to the nascent activating synapse is energy efficient and as such makes evolutionarily sense. Modulation of SHP-1 activity, its expression level, or its recruitment to the synapses could be used as therapeutic manipulation, for example in an immune therapy setting ^77^.

## MATERIALS AND METHODS

### Mice

All the animal procedures were approved by the animal ethics committee in Stockholm, Sweden (Stockholms Södra Djurförsöksetiska Nämnd and Linköpings Djurförsöksetiska Nämnd). Mice were taken care of by the animal facility at Karolinska Institute, Huddinge, Sweden. Experimental mice were at the age of six to ten weeks. For all experiments, the mice were age and gender-matched. H2K^b^D^b^ KO mice were previously generated on the C57BL/6J background ^78^ and are denoted throughout this study as Dd−/−. H2K^b^D^b^ KO mice expressing an H2D^d^ transgene were created previously and are denoted as Dd+/+ ^7, 20, 26^.

### Antibodies and flow cytometry

Antibody surface stainings were performed for 20 minutes at 4°C in FACS buffer (PBS plus 2 % FBS). Live/dead distinction was determined using eBioscience™ Fixable Viability Dye eFluor™ 780 or Aqua 405 nM (Thermofisher). Intracellular antibody staining was carried out with antibodies against IFN-g, SHP-1, SHP-2, SHIP1 (as in the antibody table) using the Foxp3/Transcription Factor Staining Buffer Kit (eBiosciences). The intracellular staining procedure was described in our other study ^79^. Data were acquired on an LSR Fortessa or a Symphony flow cytometer (BD Biosciences) and analyzed using FlowJo v9.9.6 or v10.7.1 (Tree Star). Antibodies used in this study are provided in **Supplementary table 1**. For phosphoflow experiments, enriched NK cells were stained with surface antibody and biotin-rat-anti mouse NKp46 (29A1.4, Biolegend) as described above. The stimulation, barcoding and staining were performed as described in our previous publications ^45, 80^. The barcoding dyes which were used in this study were Pacific Orange and Alexa Fluor 700 NHS esters (Invitrogen).

### Mouse and human NK cell isolation

Mouse NK cells were enriched from a single cell suspension of total splenocytes using a customized negative selection protocol as described previously ^45^. Human NK cells were enriched from peripheral blood mononuclear cells (PBMC) using the human NK cell isolation kit (Miltenyi).

### Single cell RNA sequencing

Enriched NK cells from spleens of Dd+/+ and Dd−/− mice were stained with antibodies against NK1.1, CD3, CD19, Ter-119, Ly49CI, Ly49G2, Ly49A, NKG2A and Aqua Live Dead as described above. Cells from each strain were then labeled with TotalSeq™-A0301 anti-mouse Hashtag Antibody against CD45 and MHC-I (BioLegend, Clone M1/42 and 30-F11, Hashtag 1, 2, 3) for 20 min at 4°C. The experiments were repeated three times with each time having one mouse from each strain. Cells were then sorted using BD FACSAria Fusion (BD Biosciences) to achieve more than 99 % purity. Subsequently, cells were pooled and subjected to 10x Genomics’ single-cell RNA sequencing, run by Eukaryotic Single Cell Genomics Facility, Science for Life Laboratory, Stockholm.

### scRNA-seq data analysis

All analysis was conducted in R (v. 3.6.2). Hashtag oligos were demultiplexed using HTODemux in Seurat (v. 3.1.5) with the positive quantile set to 0.99. Demultiplexing results were visually inspected, and falsely annotated cells were manually removed from the H2-D1 −/− samples. Spillover between H2-D1 +/− and H2-D1 +/+ was visually inspected and deemed to be minimal and not to affect the results. Before further analysis the data were filtered for low (<1100) and high (>4100) number of features, and mitochondrial, ribosomal, X- and Y-chromosome genes were removed. H2-D1 was likewise excluded from the analysis but kept as an independent assay to visualize the expression on UMAPs. Dimensionality reduction, UMAP visualization, and differential expression analysis were all done using Seurat (v. 3.1.5). The top 1000 variable features (determined using the vst selection method) were used for dimensionality reduction. UMAPs were generated using PCA-guided projections of log-normalized and scaled counts with the following settings: n.components=10, n.neighbors=30, n.epochs=200, min.dist=0.3, learning.rate=1, and spread=1. Differential gene expression analysis was done with FindMarkers in Seurat, with the following settings: logfc.threshold = 0, test.use = “wilcox”, min.pct = 0.1, only.pos = FALSE, min.cells.group = 10, min.cells.feature = 10, and pseudocount.use = 1. Results were visualized using ggplot2 (v. 3.3.0) and reported in Supplementary Table 1). Gene ontology analysis was performed on all significant differentially expressed genes (p value < 0.05) with clusterProfiler (v. 3.14.3) and org.Mm.eg.db (v. 3.10.0). Additionally, the following dependent package versions were installed: DOSE (v. 3.12.0), AnnotationDbi (v. 1.48.0), IRanges (v. 2.20.2), S4Vectors (v. 0.24.4), BiocGenerics (v. 0.32.0), and Biobase (v. 2.46.0). The scRNAseq generated during the current study are available in the ArrayExpress repository, under accession number: E-MTAB-XXXX (reviewer link: http://www.ebi.ac.uk/arrayexpress/experiments/E-MTAB-XXX, username: Reviewer_E-MTAB-XXXX, password: santa). All scripts used for bioinformatics analysis are available on Github: https://github.com/jonasns/NKscRNAseq.

### In-vitro stimulation experiments

#### For mouse NK cells

Enriched NK cells were stimulated with plate bound anti-NK1.1 antibody. Stimulations and stainings were performed as described previously ^81^. After 4.5-hour stimulation, cells were transferred to V-bottom plate for antibody staining. Fc block and surface antibody staining were carried out as described above. Cells were fixed, permeabilized, and stained with anti-IFN-γ using the Foxp3/Transcription Factor Staining Buffer Kit (eBiosciences).

#### For human NK cells

Flat-bottom plate were coated with 20 μg/ml AffiniPure F(ab’)_2_ Fragment Donkey Anti-Mouse IgG (H+L)mouse (Jackson Immunoresearch) in PBS for two hours at 37°C. The plates were washed twice with PBS plus 2 % FBS and subsequently coated with 10 μg/ml unconjugated anti-CD16 (3GB, BioLegend) in PBS overnight at 4°C. Enriched human NK cells were resuspended in RPMI plus 10 % FBS and anti-CD107a antibody and dispensed on the anti-CD16 coated plate. The stimulation was performed similarly to the mouse NK cell experiments.

### Quantitative PCR

NK cells were sorted as live NK1.1^+^CD3^-^ cells using BD AriaIII (BD Biosciences) FACS machine. Cells were lysed and RNA purification was performed using the RNeasy micro kit (Qiagen). cDNA synthesis was carried out using High-Capacity cDNA Reverse Transcription Kit with RNase Inhibitor (Life Technologies). Quantitative PCR (Q-PCR) was performed using standard Taqman universal mastermix and primers (Life Technologies). The primers (Life Technologies) were *ptpn6* (Mm00469153_m1), *ptpn11* (Mm00448434_m1), *inpp5d* (Mm00494987_m1), *ptprs* (Mm00465150_m1), *ppp4c* (Mm00479960_g1), *gapdh* (Mm99999915_g1).

### Sample preparation for microscopy

μ-Slide 18 Well Glass Bottom (Ibidi) plates were coated with 0.01 % poly-L-Lysine (Sigma) for 30 minutes at RT. The plates were washed three times with water and dried on a 60°C heat block. 20 μg/ml polyclonal goat anti-mouse NKp46 (R&D Systems) in PBS was then dispensed on the plate and the incubation was done for 2 hours at 37°C. The plates were washed three times using PBS and stored in PBS for up to 4 hours before the cells being added. For some experiments, enriched NK cells were used. For some experiments **(Fig. 6g, 7a-f, 8a)**, cell sorting using FACS was performed to obtain CD3-NKG2A-Ly49C-Ly49G2-LY49I-cells **(Fig. S5b)**. Sorted cells were allowed to recover for two hours in RPMI 10 % FBS at 37°C before the stimulation. A maximum of 100.000 sorted cells were then dispensed onto each well of Ibidi plates, briefly spun at 100 x g for 15 seconds and incubated for five minutes at 37°C. The cells were then fixed immediately with same volume of 4 % formaldehyde (16 % no methanol format, Thermofisher) for 20 minutes at RT. Blocking was done with Fc blocker (Innovex Biosciences) for 30 minutes at RT. Subsequently, the cells were stained with unconjugated rat anti-Ly49A antibody (YE1/32.8.5, Stemcell) followed by AF555 conjugated goat-anti-rat IgG (Life Technologies), both were done for 45 minutes in PBS at RT. The permeabilization and intracellular staining were done similarly to the flow cytometry intracellular staining with the exception that the duration of staining was 45 minutes for each step. Between each staining steps, plates were washed 5 times with either PBS (for Ly49A staining) or permeabilization buffer (for intracellular staining). Before recording the images, phalloidin conjugated with AF488, AF568 (Life Technologies), or A635P (Abberior) in PBS was added.

For experiments with resting cells, enriched NK cells were fixed with 1 % formaldehyde (diluted in PBS from 16 %, non-methanol format, Thermofisher) for 20 minutes at RT, followed by blocking with peptide-based Fc blocker (Innovex Biosciences) for 30 minutes at RT. Subsequently, cells were stained with purified anti-Ly49A antibody (YE1/32.8.5, Stem Cell) and followed by goat-anti-rat IgG antibody conjugated with Abberior STAR 635P (Abberior), both were done in PBS for 45 minutes at RT. The cells were then stained with antibodies against CD3, CD19, TER-119, Ly49C, Ly49I, NKG2A, Ly49G2, CD11b, CD27 (refer to **Supplementary table S1**). Cells that were negative for CD3, CD19, TER119, Ly49C, Ly49I, Ly49G2, NKG2A were further stratified and sorted according to CD27 and CD11b expression. The sorted populations were CD27^high^CD11^low^, CD27^high^CD11b^high^ and CD27^low^CD11b^high^ (**Fig. S6)**. Subsequently, the intracellular staining was performed similarly to the confocal experiments. Secondary AF594 goat anti-rabbit antibody was used to detect SHP-1. After staining, the cells were resuspended in PBS before cytospinning (3 minutes, 700 rpm, medium speed) onto coverslips (1.5H, NordicBiolabs). The cells were then mounted onto glass slides using ProLong™ Gold Antifade Mountant (Thermofisher).

### Confocal and Stimulated emission depletion (STED) microscopy

Confocal images were acquired on a Single point scanning confocal microscope. In brief, large fields of view were taken with 60x oil objective in three channels (Ly49A, actin and transmitted light). Cells positive for Ly49A and displayed ‘activated’ actin accumulation phenotype were chosen for recording. Nyquist images were then recorded automatically at a single plane for synapse images or z stack with 9-μm-thick volume at 0.17-μm steps with PFS (perfect focus system) being on.

STED imaging was performed with a Leica TCS SP8 STED 3X (Leica Microsystems) equipped with a HCPLAPO100x/1.40 Oil STED WHITE objective. For 2-color STED imaging, the fluorophores Alexa Fluor 594 and Abberior STAR 635P were excited by a pulsed white light laser with lines at 594 nm and 653 nm, respectively. The pulsed STED laser at 775nm was used for both imaging channels. The channels were recorded sequentially with a pixel size of 20 nm at line scan speeds of 200 Hz. Cells recorded were chosen to be Ly49A+ (635 nm). 3-color STED microscopy was achieved using fluorophores Alexa Fluor 488 depleted using 592 nm continuous-wave fibre laser and Alexa Fluor 594 and Alexa Fluor 647 depleted using 775nm pulsed white light laser. The channels were recorded sequentially with a pixel size of 28.4 nm at line scan speeds of 200 Hz. A confocal channel excited at 405 nm was also used to detect Ly49 stained with Alexa Fluor 405.

### Image analysis

Images were deconvolved using either Huygens Essential (for STED experiments) or NIS-Elements Analysis (for confocal experiments). Images were analysed using pipelines made in CellProfiler (v3.1.9). Protein colocalization analysis was done on deconvolved images. And staining protein fluorescence intensity analysis was performed on raw images. In brief, channels were splitted and each cell was identified as “primary objects” by performing segmentation on the actin channel. Subsequently, the object intensity, area occupied, and size shape were measured. Ly49A images with numbered primary objects were exported and used to select Ly49A+ cells manually. For the analysis of synaptic regions, secondary objects were identified by shrinking primary objects by 10 pixels (1.13 μm). The subtraction of primary objects to secondary objects was taken as dSMAC. Similarly, tertiary objects (cSMAC) were identified by shrinking secondary objects by 10 pixels (1.13 μm). The subtraction of secondary objects to tertiary objects was taken as pSMAC. Subsequently, the intensity of different channels and the areas of dSMAC, pSMAC, and cSMAC were measured. The data presented in **Fig. 6d,e,g** were the intensity per region normalized to the average of Dd−/− dSMAC in each experiment. Object-based distance analysis was also done using a CellProfiler pipeline. A gaussian filter and an enhance filter were applied to reduce the background signal. Clusters of SHP-1 were identified as primary objects and images of the primary objects were then created. After these images were reverted, a distance operation was applied. In the end, SLP76 (pY128) objects were placed on these images and received distance values, which reflected the distance to surrounding SHP-1 objects. As SLP76 (pY128) objects might have taken several distance values, the minimum distance was plotted, indicating the shortest distance to the surrounding SHP-1 objects. Thereafter, the average of these distances per cell or for all objects in each cell were plotted. A similar analysis for SLP76 objects to surrounding SHP-1 (pY564) objects was performed.

Intensity analysis from z stacks was done automatically with Fiji Macro and the files were combined using R. The slice with the highest actin intensity was taken as an anchor. Two-μm volume around the highest actin-intensity slice was designated as the synapse area. SHP-1 intensity in each slice was subtracted to the lowest background intensity among all the images of the first slice from the bottom (near the synapse). The ratio of sum SHP-1 intensity in the synapse area over that in the whole cell was plotted as the SHP-1 enrichment in the synapse area.

### CRISPR-cas9 gene modulation

#### Common

All CRISPR RNA (crRNA) and tracer RNA (trcRNA) were ordered from Integrated DNA Technologies (IDT) in their proprietary Alt-R format (www.idtdna.com/CRISPR-Cas9). SgRNA and trcRNA were reconstituted at 200 μM in Nuclease-Free Duplex Buffer (IDT). For each reaction, Cas9 recombinant protein (1.8 μl of 60 μM concentration equals to 108 pmol, QB3 lab, Berkeley lab) was added to the sgRNA or trcRNA for control (1.2 μl equals 120 pmol). The mixture was incubated for ten-minute incubation at RT to form Crispr-Cas-ribonucleoprotein (RNP). Enriched NK cells (mouse or human) were resuspended in a custom-made electroporation buffer (5 mM KCl, 15 mM MgCl2, 120 mM Na_2_HPO4/NaH_2_PO_4_ (pH7.2), 50 mM Manitol) at the concentration of a maximum of one million cells per 20 μl buffer per reaction. Twenty μl of cells in buffer were transferred to each well of a 16-well Nucleofection cuvette strip (4D-Nucleofector X kit S; Lonza). Three μl of RNP were gently dispensed to each well, avoiding bubbles. The electroporation was done using a 4D nucleofector (4D-Nucleofector Core Unit: Lonza, AAF-1002B; 4D-Nucleofector X Unit: AAF-1002X; Lonza), with CM137 as the pulse program and P3 as the selected electroporation buffer. Eighty μl of pre-warmed RPMI plus 10 % FBS were added onto each well of the Nucleofection strip. The cell solution was transferred to a U-bottom 96-well plate which contained 100 μl pre-warmed RPMI plus 10 % FBS and two-time desired concentration of cytokines (indicated below for either mouse or human cells). The cells were incubated with the indicated number of days with the replacement of 75 μl RPMI plus 10 % FBS and cytokines every day. Before downstream functional assays, the cells were washed twice to remove cytokines.

#### For mouse NK cells

NK cells were enriched and incubated with 50 ng/ml recombinant mouse IL-15 (Preprotech) for four days in RPMI plus 10 % FBS with penicillin and streptomycin (Thermofisher) added in a U-bottom 96-well plate at the concentration of two million cells per ml. Half of the medium was replaced with fresh RPMI plus 10 % FBS and 10 ng/ml IL-15 every day. Following electroporation, the cells were incubated with RPMI plus 10 % FBS without any antibiotic added and 10 ng/ml IL-15 for at least three days. The cells were washed twice to remove IL-15 before dispensed on anti-NK1.1 coated 96-well flat-bottom plate. Microscopy experiment, calcium flux assay, SHP-1 staining, IFN-γ production and degranulation assay was carried as described above.

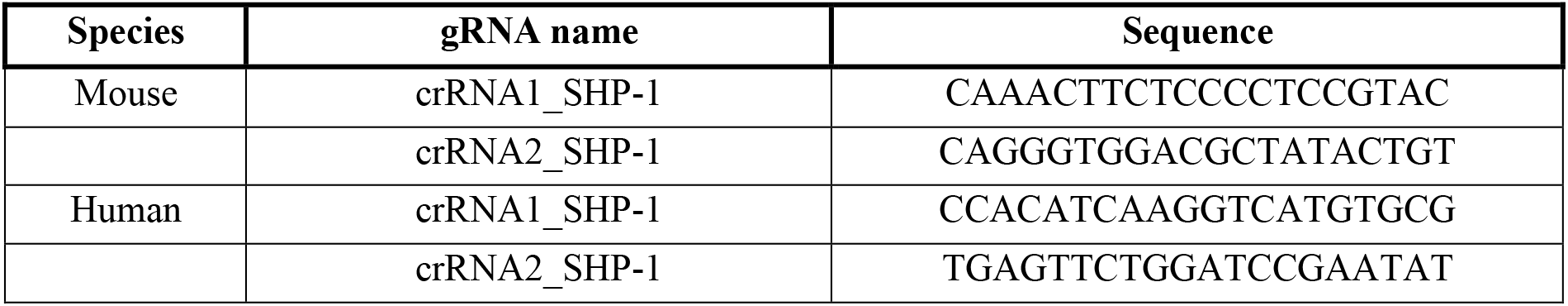

#### For human NK cells

A similar protocol was performed for human NK cells with the exceptions that RPMI plus 10 % FBS and penicillin and streptomycin (Sigma) and 5 ng/ml recombinant human IL-15 (Preprotech) were used.

### SHP-1 inhibition

#### For mouse NK cells

Enriched mouse NK cells from Dd−/− and Dd+/+ mice were incubated with SHP-1 inhibitor (denoted in the paper as iSHP-1) (NSC-87877, CAS 56932-43-5 – Calbiochem) of different concentrations or PBS of the amount corresponding to the highest iSHP-1 concentration. The incubation was done overnight with 2 ng/ml recombinant-mouse IL-15 (Preprotech) in RPMI plus 10 % FBS. After the overnight incubation (about 20 hours), NK cells together with iSHP-1 and IL-15 were transferred to an anti-NK1.1 coated plate and the assay was carried out as described above for the mouse plate-bound antibody experiments. After 4.5-hour stimulation, the cells were washed 3 times in ice-cold PBS plus 2 % FBS to remove completely iSHP-1 (the compound was colorful). Subsequently, the cells were labeled with surface antibodies, fixed, permeabilized, and stained with anti-IFN-g antibody as described above.

#### For human NK cells

Enriched human NK cells were incubated with iSHP1 (same as for mouse NK cell experiment) of different concentrations or DMSO of the amount corresponding to the highest iSHP1 concentration. The incubation was done for 2.5 hours in RPMI plus 10 % FBS. Subsequently, NK cells together with iSHP-1 were transferred to an anti-CD16 coated plate and the assay was carried out as described above for the human plate-bound antibody experiments. After 4.5-hour stimulation, the cells were washed 3 times in ice-cold PBS plus 2 % FBS to remove completely iSHP-1. NK cells were stained with surface antibodies, fixed, permeabilized and stained with anti-IFN-g antibody as described above.

### Statistical analysis

Data were analysed and plotted with Graphpad Prism version 6 for Mac OSX (GraphPad Software, La Jolla California USA, www.graphpad.com).

## Acknowledgements

We thank the PKL4 animal facility (Karolinska Institutet, Huddinge) staff for taking care of the mice. We would like to acknowledge Iyadh Douagi, Belinda Pannagel, Mahin Nikougoftar Zarif, and the MedH Flow Cytometry core facility, supported by a joint core facility grant from the Karolinska Insitutet and the Stockholm City Council, for providing cell analysis and sorting services. Single-cell transcriptomic data were generated at the Eukaryotic Single-Cell Genomics facility at Science for Life Laboratory (SciLife Lab) in Stockholm, Sweden. Data storage was supported by the Swedish National Infrastructure for Computing (SNIC) at the UPPMAX. We also thank the Centre for Hematology and Regenerative Medicine (HERM), Karolinska Institute, for providing a great facility and scientific environment. We also acknowledge Hans Blom, Daniel Dans and the Advanced Light Microscopy facility at SciLife Lab, Stockholm, Sweden of the National Microscopy Infrastructure, NMI (VR-RFI 2019-00217) for assistance with STED microscopy.

## Funding

This work was funded by grants to P.H. from the Swedish Research Council, the Swedish Cancer Society, Region Stockholm, Radiumhemmets Forskningsfonder, Aroseniusfonden and Karolinska Institutet. T.T.L. was supported by a KID grant from Karolinska Institutet. LS was supported by the Swiss National Science Foundation (SNF): P400PM_183909.

## Author contributions

L.S. and T.T.L. conceived the project, performed experiments, analysed data and wrote the manuscript. P.H. conceived the project and wrote the manuscript. J.N.S. analyzed scRNA-seq data. M.B.S. contributed to STED experiments, provided methodology, scientific inputs, and manuscript edits. D.K.M., S.B.S., C.M., H. M. T, G.P.C., A.K.W., G.R., G.G. performed experiments and manuscript edits. M.C., H.S., L.W., Y.B., E.A., N.K., S.M. contributed to method optimization and manuscript edits.

## Competing interests

The authors declare that they have no competing interests.

## SUPPLEMENTARY FIGURE LEGENDS

**Fig. S1. Gating strategy for mouse NK cell subsets and scRNA-seq sorting. a**, Gating strategy to define sp subsets of murine NK cells. Boolean gating was used to analyze 32-subsets expressing a combination of five main inhibitory receptors in B6 mice. **b**, Gating strategy for FACS sorting of sp-Ly49A for scRNA-seq. **c,** Pipeline for the scRNA-seq experiment. Enriched NK cells were stained with oligo-labelled antibody prior to FACS sorting to isolate sp-Ly49A subsets. Cells from Dd+/+ and Dd−/− mice were pooled and subjected to 10x Genomics RNA scRNA-seq. Three independent experiments with one mouse from each strain per experiment were performed. **d**, H-2D1 (Dd) expression before and after filtering. **e,** UMAPs of selected genes. **f,** Pathway enrichment analysis for biological processes (BP) of the differentially expressed genes (DEGs), FDR<0.05.

**Fig. S2. SHP-1 staining of mouse NK cells. a,** mRNA abundance in NK cells of *ptpn6, ptpn11* and *inpp5d* as compared to *β-actin* and *gapdh*. **b,** Control staining for C14H6 antibody (CST) which was used throughout the paper to detect mouse SHP-1. Flow cytometry was used to detect SHP-1 intracellularly from CD4+ cells from spleens of CD4^Cre-/-^ Ptpn6^flox/flox^ (controls) and CD4^Cre+/+^ Ptpn6^flox/flox^ mice. **c**, Gating strategy to define CD11b+ and CD11b-populations. Cells were labelled with cell trace violet (CTV) prior to surface antibody staining. **d,** SHP-1 expression on the three maturation subsets segregated by CD27 and CD11b expression. **e, f**, SHP-2 and SHIP1 expression on CD11b- and CD11b+ NK populations from Dd−/− and Dd+/+ mice. **d, e, f,** Two-way ANOVA with Sidak’s multiple comparisons. *p < 0.05, **p < 0.01, ***p < 0.001.

**Fig. S3. Confirmation of human SHP-1 antibody using Crispr/Cas9 and gating strategy to stratify human NK cell subsets. a**, Peripheral NK cells from healthy donors were cultured for 3 days in the presence of 5 ng/ml IL-15 prior to electroporation. Three-day post electroporation, SHP-1 expression was measured intracellularly using W17240D antibody (Biolegend) and flow cytometry. **b**, Gating strategy for the identification of human NK cell subsets. Boolean gating was done to stratify subsets based on the expression of KIR2DL1, KIR3DL1, KIR2DL2/3 and NKG2A. Functions and MFI of SHP-1 were then measured on these subsets.

**Fig. S4. Gating strategy for phosphoflow experiments and increment ratios of phosphorylation upon NKp46 stimulation. a**, Gating strategy to identify cells from the phosphoflow experiment. NKp46 was crosslinked using biotin-conjugated NKp46 antibody and streptavidin for 60 seconds. The cells were then fixed, permeabilized and barcoded with ester amine dyes (Alexa Flour 700 and Pacific Orange). Subsequently, cells were pooled for downstream staining with phospho-antibodies. **b**, Fold increase of phosphorylation of investigated molecules upon NKp46 stimulation as compared to untreated samples.

**Fig. S5. Controls and gating strategy for microscopy experiments. a**, Controls showing specific stainings obtained by confocal microscopy. AF647 was used for SHP-1. AF555 was used for Ly49A. AF405 was used for SLP76 (pY128). An example of how Ly49A+ cells is shown. **b**, Gating strategy for the sorting experiments to get sp-Ly49A and Null subsets prior to NKp46 stimulation and staining.

**Fig. S6. Gating strategy for cell sorting for STED experiments in Fig. 8.**

## Notes

### Competing Interest Statement

The authors have declared no competing interest.

## REFERENCES

1. Long, E.O., Kim, H.S., Liu, D., Peterson, M.E. & Rajagopalan, S. Controlling natural killer cell responses: integration of signals for activation and inhibition. Annu Rev Immunol 31, 227–258 (2013).

2. Hoglund, P. & Brodin, P. Current perspectives of natural killer cell education by MHC class I molecules. Nat Rev Immunol 10, 724–734 (2010).

3. Kennedy, P.R. et al. Genetic diversity affects the nanoscale membrane organization and signaling of natural killer cell receptors. Sci Signal 12 (2019).

4. Alari-Pahissa, E., Grandclement, C., Jeevan-Raj, B. & Held, W. Inhibitory receptor-mediated regulation of natural killer cells. Crit Rev Immunol 34, 455–465 (2014).

5. Raulet, D.H. & Guerra, N. Oncogenic stress sensed by the immune system: role of natural killer cell receptors. Nat Rev Immunol 9, 568–580 (2009).

6. Kadri, N., Thanh, T.L. & Hoglund, P. Selection, tuning, and adaptation in mouse NK cell education. Immunol Rev 267, 167–177 (2015).

7. Johansson, S. et al. Natural killer cell education in mice with single or multiple major histocompatibility complex class I molecules. J Exp Med 201, 1145–1155 (2005).

8. Kim, S. et al. Licensing of natural killer cells by host major histocompatibility complex class I molecules. Nature 436, 709–713 (2005).

9. Anfossi, N. et al. Human NK cell education by inhibitory receptors for MHC class I. Immunity 25, 331–342 (2006).

10. Boudreau, J.E. & Hsu, K.C. Natural killer cell education in human health and disease. Curr Opin Immunol 50, 102–111 (2018).

11. Hoglund, P. et al. Natural resistance against lymphoma grafts conveyed by H-2Dd transgene to C57BL mice. J Exp Med 168, 1469–1474 (1988).

12. Bix, M. et al. Rejection of class I MHC-deficient haemopoietic cells by irradiated MHC-matched mice. Nature 349, 329–331 (1991).

13. Fernandez, N.C. et al. A subset of natural killer cells achieves self-tolerance without expressing inhibitory receptors specific for self-MHC molecules. Blood 105, 4416–4423 (2005).

14. Hoglund, P. et al. Recognition of beta 2-microglobulin-negative (beta 2m-) T-cell blasts by natural killer cells from normal but not from beta 2m-mice: nonresponsiveness controlled by beta 2m-bone marrow in chimeric mice. Proc Natl Acad Sci U S A 88, 10332–10336 (1991).

15. Zimmer, J. et al. Phenotypic studies of natural killer cell subsets in human transporter associated with antigen processing deficiency. PLoS One 2, e1033 (2007).

16. Zhang, X., Feng, J., Chen, S., Yang, H. & Dong, Z. Synergized regulation of NK cell education by NKG2A and specific Ly49 family members. Nat Commun 10, 5010 (2019).

17. Raulet, D.H. & Vance, R.E. Self-tolerance of natural killer cells. Nat Rev Immunol 6, 520–531 (2006).

18. Chen, S. et al. The Self-Specific Activation Receptor SLAM Family Is Critical for NK Cell Education. Immunity 45, 292–304 (2016).

19. Wu, N. et al. A hematopoietic cell-driven mechanism involving SLAMF6 receptor, SAP adaptors and SHP-1 phosphatase regulates NK cell education. Nat Immunol 17, 387–396 (2016).

20. Brodin, P., Lakshmikanth, T., Johansson, S., Karre, K. & Hoglund, P. The strength of inhibitory input during education quantitatively tunes the functional responsiveness of individual natural killer cells. Blood 113, 2434–2441 (2009).

21. Brodin, P., Karre, K. & Hoglund, P. NK cell education: not an on-off switch but a tunable rheostat. Trends Immunol 30, 143–149 (2009).

22. Brodin, P. & Hoglund, P. Beyond licensing and disarming: a quantitative view on NK-cell education. Eur J Immunol 38, 2934–2937 (2008).

23. Joncker, N.T., Fernandez, N.C., Treiner, E., Vivier, E. & Raulet, D.H. NK cell responsiveness is tuned commensurate with the number of inhibitory receptors for self-MHC class I: the rheostat model. J Immunol 182, 4572–4580 (2009).

24. Wu, Z. et al. Dynamic variability in SHP-1 abundance determines natural killer cell responsiveness. Sci Signal 14, eabe5380 (2021).

25. Boudreau, J.E. et al. Cell-Extrinsic MHC Class I Molecule Engagement Augments Human NK Cell Education Programmed by Cell-Intrinsic MHC Class I. Immunity 45, 280–291 (2016).

26. Brodin, P., Lakshmikanth, T., Karre, K. & Hoglund, P. Skewing of the NK cell repertoire by MHC class I via quantitatively controlled enrichment and contraction of specific Ly49 subsets. J Immunol 188, 2218–2226 (2012).

27. Boos, M.D., Yokota, Y., Eberl, G. & Kee, B.L. Mature natural killer cell and lymphoid tissue-inducing cell development requires Id2-mediated suppression of E protein activity. J Exp Med 204, 1119–1130 (2007).

28. Muller-Durovic, B. et al. Killer Cell Lectin-like Receptor G1 Inhibits NK Cell Function through Activation of Adenosine 5’-Monophosphate-Activated Protein Kinase. J Immunol 197, 2891–2899 (2016).

29. Viant, C. et al. SHP-1-mediated inhibitory signals promote responsiveness and anti-tumour functions of natural killer cells. Nat Commun 5, 5108 (2014).

30. Maghazachi, A.A., Al-Aoukaty, A. & Schall, T.J. CC chemokines induce the generation of killer cells from CD56+ cells. Eur J Immunol 26, 315–319 (1996).

31. Awasthi, A. et al. Rap1b facilitates NK cell functions via IQGAP1-mediated signalosomes. J Exp Med 207, 1923–1938 (2010).

32. Crinier, A. et al. High-Dimensional Single-Cell Analysis Identifies Organ-Specific Signatures and Conserved NK Cell Subsets in Humans and Mice. Immunity 49, 971–986 e975 (2018).

33. Yang, C. et al. Single-cell transcriptome reveals the novel role of T-bet in suppressing the immature NK gene signature. Elife 9 (2020).

34. Wagner, A.K. et al. Retuning of Mouse NK Cells after Interference with MHC Class I Sensing Adjusts Self-Tolerance but Preserves Anticancer Response. Cancer Immunol Res 4, 113–123 (2016).

35. Chiossone, L. et al. Maturation of mouse NK cells is a 4-stage developmental program. Blood 113, 5488–5496 (2009).

36. Kim, S. et al. In vivo developmental stages in murine natural killer cell maturation. Nat Immunol 3, 523–528 (2002).

37. Chen, L. et al. Discovery of a novel shp2 protein tyrosine phosphatase inhibitor. Mol Pharmacol 70, 562–570 (2006).

38. Beziat, V. et al. NK cell responses to cytomegalovirus infection lead to stable imprints in the human KIR repertoire and involve activating KIRs. Blood 121, 2678–2688 (2013).

39. Corral, L., Hanke, T., Vance, R.E., Cado, D. & Raulet, D.H. NK cell expression of the killer cell lectin-like receptor G1 (KLRG1), the mouse homolog of MAFA, is modulated by MHC class I molecules. Eur J Immunol 30, 920–930 (2000).

40. Bjorkstrom, N.K. et al. Expression patterns of NKG2A, KIR, and CD57 define a process of CD56dim NK-cell differentiation uncoupled from NK-cell education. Blood 116, 3853–3864 (2010).

41. Stebbins, C.C. et al. Vav1 dephosphorylation by the tyrosine phosphatase SHP-1 as a mechanism for inhibition of cellular cytotoxicity. Mol Cell Biol 23, 6291–6299 (2003).

42. Matalon, O. et al. Dephosphorylation of the adaptor LAT and phospholipase C-gamma by SHP-1 inhibits natural killer cell cytotoxicity. Sci Signal 9, ra54 (2016).

43. Binstadt, B.A. et al. SLP-76 is a direct substrate of SHP-1 recruited to killer cell inhibitory receptors. J Biol Chem 273, 27518–27523 (1998).

44. Plas, D.R. et al. Direct regulation of ZAP-70 by SHP-1 in T cell antigen receptor signaling. Science 272, 1173–1176 (1996).

45. Luu, T.T. et al. Short-term IL-15 priming leaves a long-lasting signalling imprint in mouse NK cells independently of a metabolic switch. Life Sci Alliance 4 (2021).

46. Kosugi, A., Sakakura, J., Yasuda, K., Ogata, M. & Hamaoka, T. Involvement of SHP-1 tyrosine phosphatase in TCR-mediated signaling pathways in lipid rafts. Immunity 14, 669–680 (2001).

47. Wang, W. et al. Crystal structure of human protein tyrosine phosphatase SHP-1 in the open conformation. J Cell Biochem 112, 2062–2071 (2011).

48. Yang, J. et al. Crystal structure of human protein-tyrosine phosphatase SHP-1. J Biol Chem 278, 6516–6520 (2003).

49. Yoshida, K., Kharbanda, S. & Kufe, D. Functional interaction between SHPTP1 and the Lyn tyrosine kinase in the apoptotic response to DNA damage. J Biol Chem 274, 34663–34668 (1999).

50. Matalon, O. et al. Actin retrograde flow controls natural killer cell response by regulating the conformation state of SHP-1. EMBO J 37 (2018).

51. Oszmiana, A. et al. The Size of Activating and Inhibitory Killer Ig-like Receptor Nanoclusters Is Controlled by the Transmembrane Sequence and Affects Signaling. Cell Rep 15, 1957–1972 (2016).

52. Vyas, Y.M. et al. Spatial organization of signal transduction molecules in the NK cell immune synapses during MHC class I-regulated noncytolytic and cytolytic interactions. J Immunol 167, 4358–4367 (2001).

53. Hammer, J.A., Wang, J.C., Saeed, M. & Pedrosa, A.T. Origin, Organization, Dynamics, and Function of Actin and Actomyosin Networks at the T Cell Immunological Synapse. Annu Rev Immunol 37, 201–224 (2019).

54. Kumari, S. et al. Actin foci facilitate activation of the phospholipase C-gamma in primary T lymphocytes via the WASP pathway. Elife 4 (2015).

55. Chalifour, A. et al. A Role for cis Interaction between the Inhibitory Ly49A receptor and MHC class I for natural killer cell education. Immunity 30, 337–347 (2009).

56. Liu, D. et al. Integrin-dependent organization and bidirectional vesicular traffic at cytotoxic immune synapses. Immunity 31, 99–109 (2009).

57. Davis, D.M. et al. The human natural killer cell immune synapse. Proc Natl Acad Sci U S A 96, 15062–15067 (1999).

58. Ben-Shmuel, A., Sabag, B., Biber, G. & Barda-Saad, M. The Role of the Cytoskeleton in Regulating the Natural Killer Cell Immune Response in Health and Disease: From Signaling Dynamics to Function. Front Cell Dev Biol 9, 609532 (2021).

59. Brown, A.C. et al. Remodelling of cortical actin where lytic granules dock at natural killer cell immune synapses revealed by super-resolution microscopy. PLoS Biol 9, e1001152 (2011).

60. Banerjee, P.P. & Orange, J.S. Quantitative measurement of F-actin accumulation at the NK cell immunological synapse. J Immunol Methods 355, 1–13 (2010).

61. Chiang, G.G. & Sefton, B.M. Specific dephosphorylation of the Lck tyrosine protein kinase at Tyr-394 by the SHP-1 protein-tyrosine phosphatase. J Biol Chem 276, 23173–23178 (2001).

62. Held, W. & Mariuzza, R.A. Cis interactions of immunoreceptors with MHC and non-MHC ligands. Nat Rev Immunol 8, 269–278 (2008).

63. Scarpellino, L. et al. Interactions of Ly49 family receptors with MHC class I ligands in trans and cis. J Immunol 178, 1277–1284 (2007).

64. Andersson, K.E., Williams, G.S., Davis, D.M. & Hoglund, P. Quantifying the reduction in accessibility of the inhibitory NK cell receptor Ly49A caused by binding MHC class I proteins in cis. Eur J Immunol 37, 516–527 (2007).

65. Back, J., Angelov, G.S., Mariuzza, R.A. & Held, W. The interaction with H-2D(d) in cis is associated with a conformational change in the Ly49A NK cell receptor. Front Immunol 2, 55 (2011).

66. Back, J., Chalifour, A., Scarpellino, L. & Held, W. Stable masking by H-2Dd cis ligand limits Ly49A relocalization to the site of NK cell/target cell contact. Proc Natl Acad Sci U S A 104, 3978–3983 (2007).

67. Coudert, J.D. et al. Altered NKG2D function in NK cells induced by chronic exposure to NKG2D ligand-expressing tumor cells. Blood 106, 1711–1717 (2005).

68. Goodridge, J.P. et al. Remodeling of secretory lysosomes during education tunes functional potential in NK cells. Nat Commun 10, 514 (2019).

69. Lowin-Kropf, B., Kunz, B., Beermann, F. & Held, W. Impaired natural killing of MHC class I-deficient targets by NK cells expressing a catalytically inactive form of SHP-1. J Immunol 165, 1314–1321 (2000).

70. Mahmood, S., Kanwar, N., Tran, J., Zhang, M.L. & Kung, S.K. SHP-1 phosphatase is a critical regulator in preventing natural killer cell self-killing. PLoS One 7, e44244 (2012).

71. Ben-Shmuel, A., Biber, G., Sabag, B. & Barda-Saad, M. Modulation of the intracellular inhibitory checkpoint SHP-1 enhances the antitumor activity of engineered NK cells. Cell Mol Immunol (2020).

72. Sosale, N.G. et al. Cell rigidity and shape override CD47’s “self-signaling in phagocytosis by hyperactivating myosin-II. Blood 125, 542–552 (2015).

73. Lee, K.H. et al. The immunological synapse balances T cell receptor signaling and degradation. Science 302, 1218–1222 (2003).

74. Oppenheim, D.E. et al. Sustained localized expression of ligand for the activating NKG2D receptor impairs natural cytotoxicity in vivo and reduces tumor immunosurveillance. Nat Immunol 6, 928–937 (2005).

75. Tripathy, S.K. et al. Continuous engagement of a self-specific activation receptor induces NK cell tolerance. J Exp Med 205, 1829–1841 (2008).

76. Sun, J.C. & Lanier, L.L. Tolerance of NK cells encountering their viral ligand during development. J Exp Med 205, 1819–1828 (2008).

77. Fernandes, R.A. et al. Immune receptor inhibition through enforced phosphatase recruitment. Nature 586, 779–784 (2020).

78. Perarnau, B. et al. Single H2Kb, H2Db and double H2KbDb knockout mice: peripheral CD8+ T cell repertoire and anti-lymphocytic choriomeningitis virus cytolytic responses. Eur J Immunol 29, 1243–1252 (1999).

79. Luu, T.T. et al. IL-15 and CD155 expression regulate LAT expression in murine DNAM1(+) NK cells, enhancing their effectors functions. Eur J Immunol (2019).

80. Schlums, H. et al. Cytomegalovirus infection drives adaptive epigenetic diversification of NK cells with altered signaling and effector function. Immunity 42, 443–456 (2015).

81. Luu, T.T. et al. Independent control of natural killer cell responsiveness and homeostasis at steady-state by CD11c+ dendritic cells. Sci Rep 6, 37996 (2016).

