## Supplementary figures and images for "Control of NK cell tolerance in MHC class I-deficiency by regulated SHP-1 localization to the activating immune synapse"

### Supplementary figures

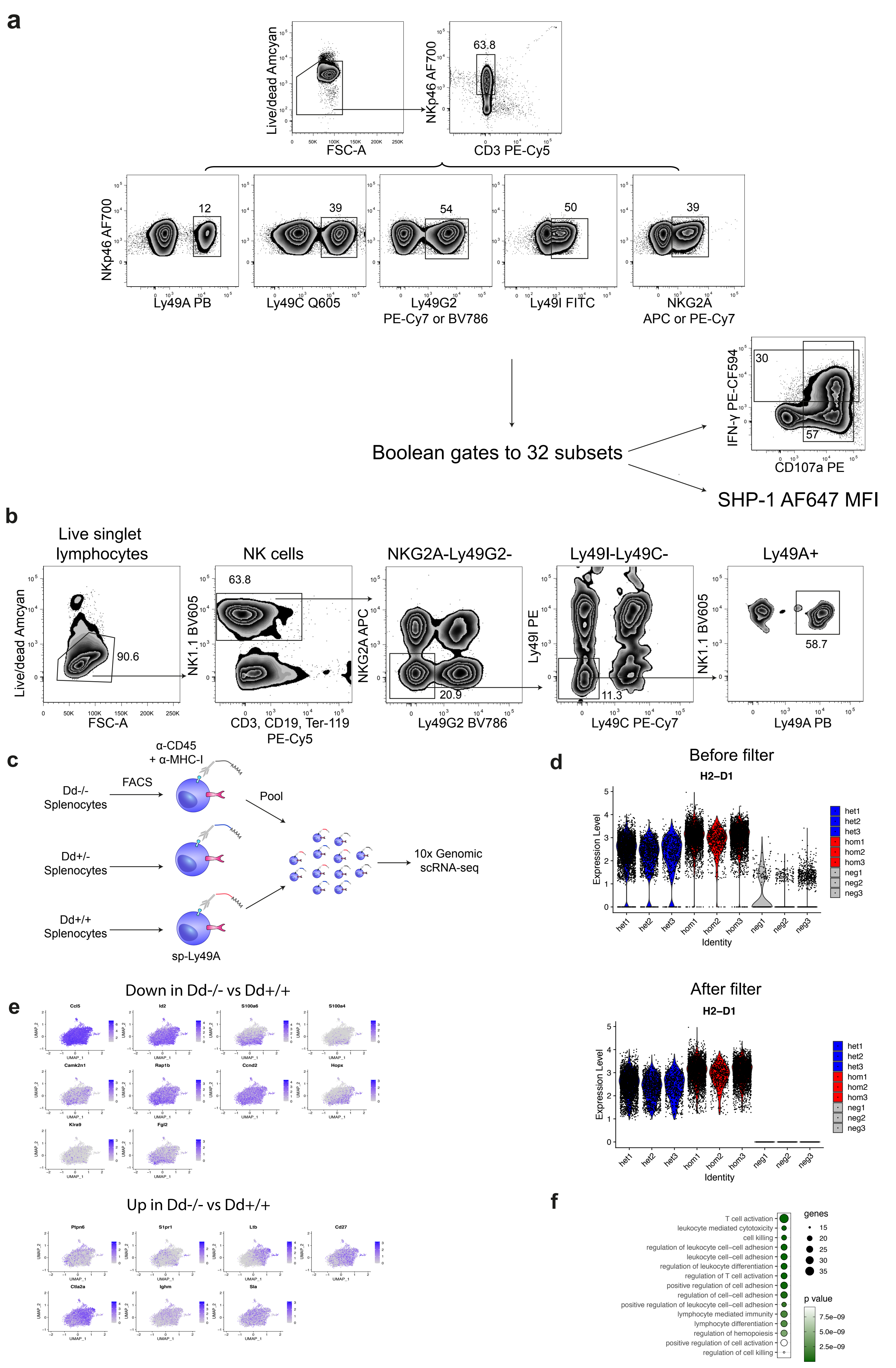

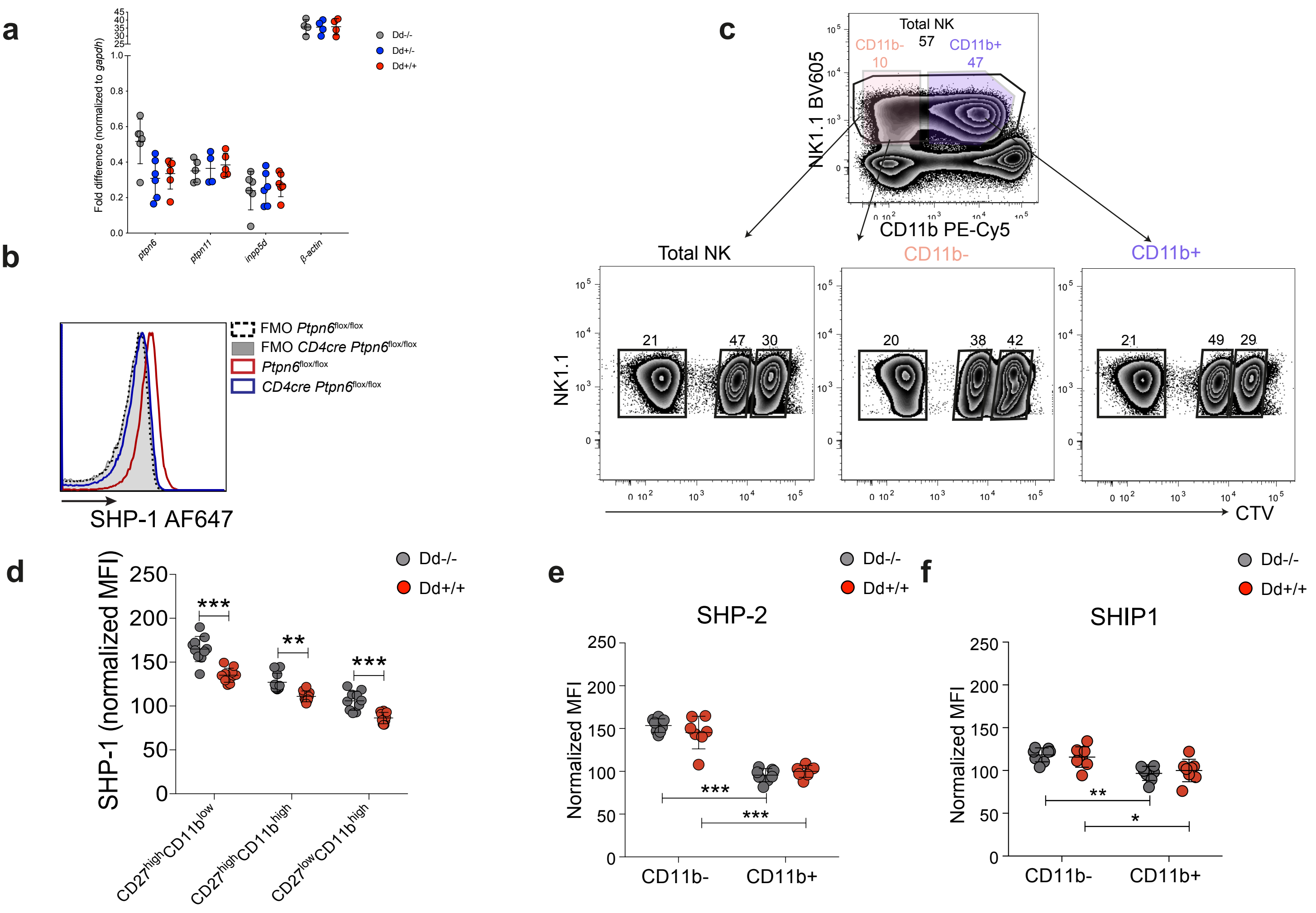

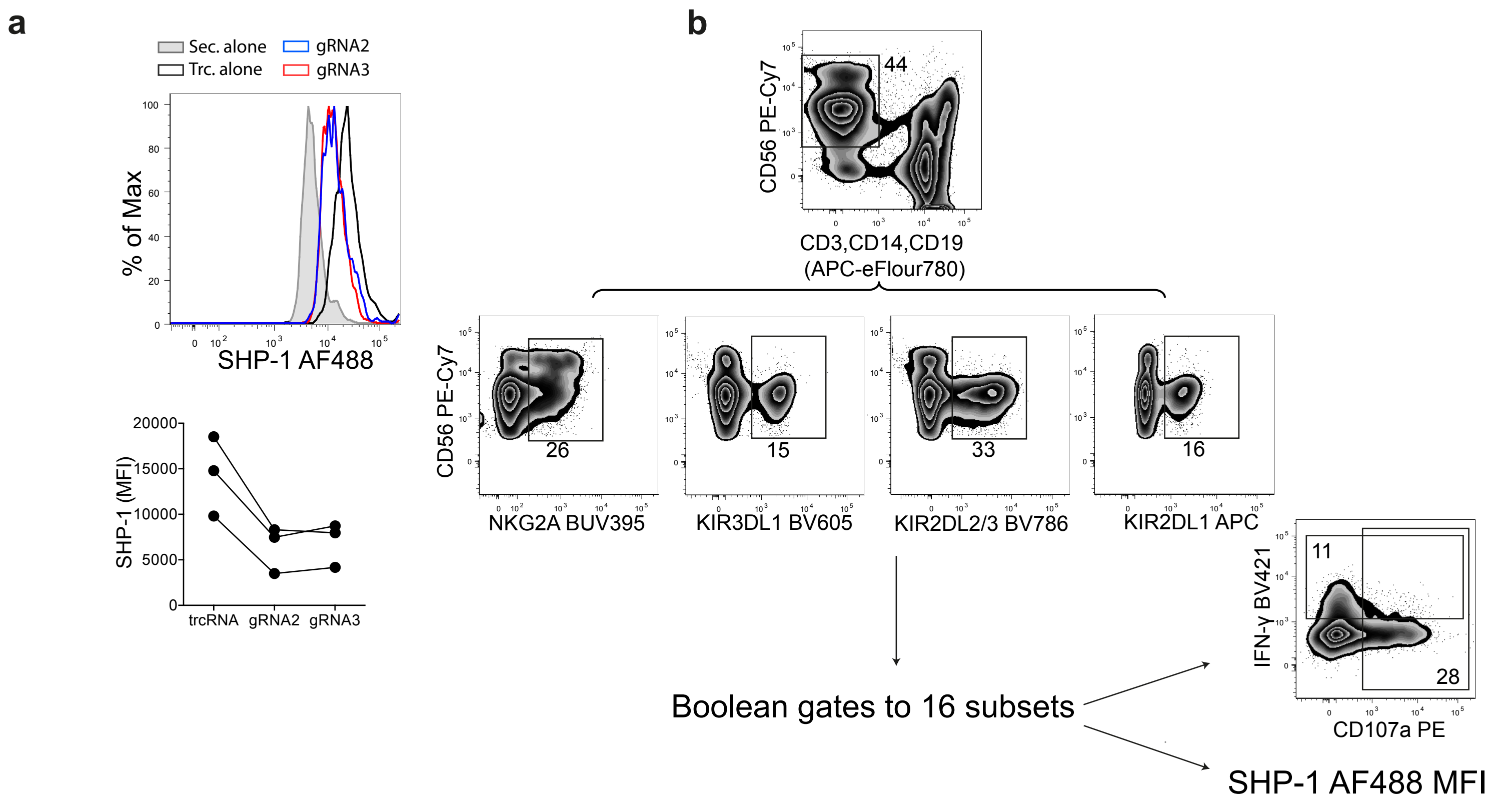

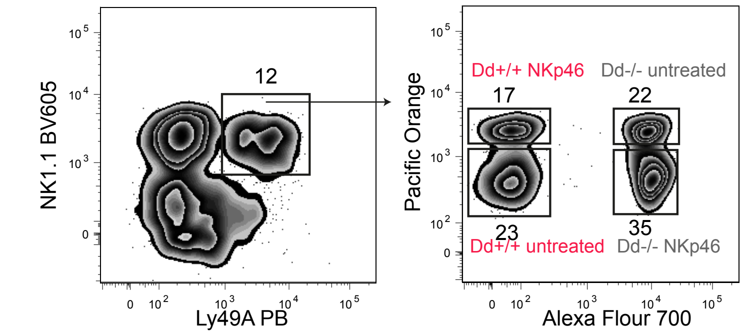

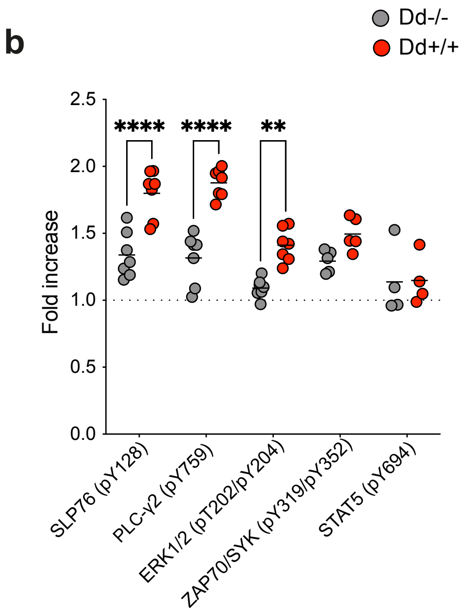

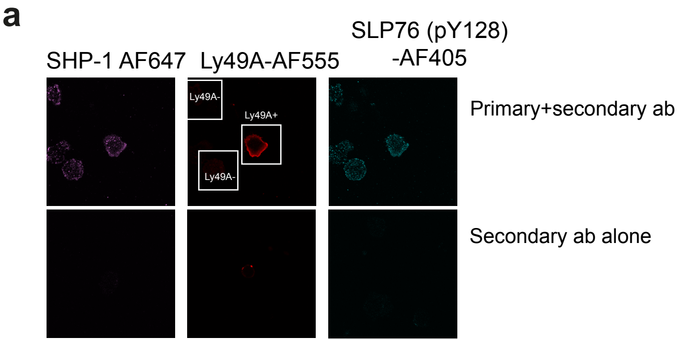

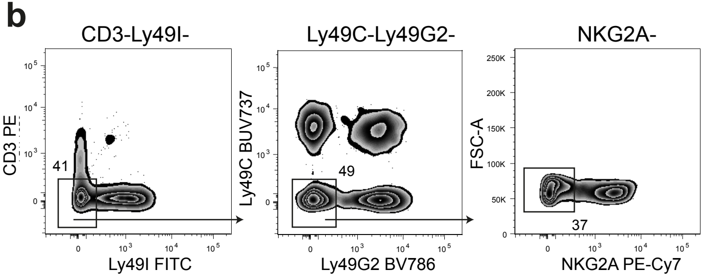

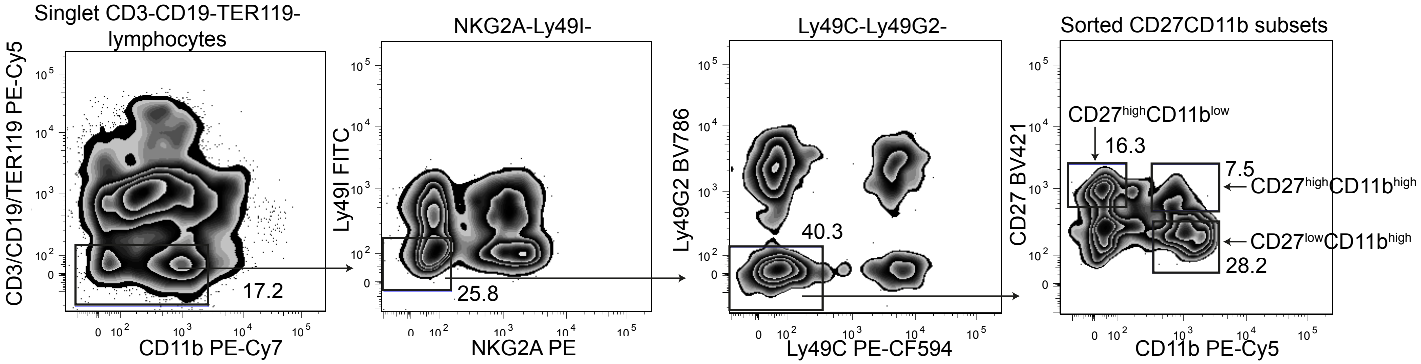
